# Membrane vesicles of *Shewanella oneidensis* MR-1 enhance denitrification growth in a species-specific manner

**DOI:** 10.1101/2025.02.19.636751

**Authors:** Kohei Takahashi, Riku Takeda, Thomas Kouyou Savage, Masahito Kataoka, Tatsuya Yamamoto, Satoshi Okabe, Nobuhiko Nomura, Mamoru Oshiki, Masanori Toyofuku, Yoshihide Tokunou

## Abstract

Denitrification, a fundamental bacterial respiratory process that occurs in anoxic environments, plays a pivotal role in energy synthesis and the global nitrogen cycle. Although the significance of this process is well-recognized, its regulation within polymicrobial communities remains poorly understood, particularly concerning interspecies interactions. In this study, we investigated the role that bacterial membrane vesicles (MV) play in modulating denitrification across bacterial species. MV is known to carry specific biomolecules such as secondary metabolites, proteins, and nucleic acids, therefore considered to be a secretion pathway. We found that MV produced by *Shewanella oneidensis* enhanced denitrification in a species-specific manner. Bacteria with highly hydrophobic surfaces tended to respond to denitrification enhancement, suggesting that the MV–bacteria attachment process is the key to generating species specificity. Transcriptome analysis and isotopic metabolite tracking indicated that the MV influenced denitrifying activities, rather than the transcription of denitrification-related genes. We further demonstrated that *c*-type cytochromes in MV act as key components that enhance denitrification. These insights expand our understanding of bacterial ecology, highlighting the role of membrane vesicles in facilitating respiratory competition and cooperation in polymicrobial communities.

## Introduction

Respiration is a fundamental energy conversion system in bacteria, where sequential redox reactions drive ATP synthesis through proton/sodium motive forces. Owing to the ability of respiration to convert substrates into reactants with different redox states, it also contributes to elemental cycling. Denitrification is a well-known process that sequentially converts nitrate (NO_3_ ^−^) to nitrite (NO_2_ ^−^), nitric oxide (NO), nitrous oxide (N_2_O), and finally to nitrogen gas (N_2_), which is an important aspect of bacterial respiration from the viewpoints of energy synthesis in anoxic environments and the nitrogen cycle [1, 2]. However, the mechanisms by which denitrification is regulated in polymicrobial communities remain poorly understood. Given that diverse bacterial species coexist in polymicrobial communities, it is plausible that various interspecies interactions are important for maintaining denitrification activity [3, 4]. These interactions may involve the sharing of metabolites, transfer of signalling molecules (e.g., quorum sensing signals), and exchange of electrons for respiration [5–8]. Certain factors within the polymicrobial community could play crucial roles in regulating bacterial respiration.

*Shewanella* thrives in environments where denitrification occurs, such as lakes and wastewater treatment plants [9, 10]. As a versatile facultative anaerobe, *S. oneidensis* has been well-studied for its ability to respire in both oxic and anoxic environments, where it utilizes a variety of electron acceptors, such as oxygen, NO_3_^−^, and metal oxides. This bacterium plays a crucial role in these communities by interacting with various bacterial species through multiple mechanisms, including interspecies electron transfer and metabolic cooperation [11–15]. Recent studies have shown that *S. oneidensis* enhances denitrification activity within microbial communities by facilitating interspecies electron transfer [16–18]. Two types of interspecies electron transfer mechanisms have been proposed to explain how *S. oneidensis* enhances denitrification activity: 1) direct electron transfer, in which physical connections via extracellular extensions, such as nanotubes and nanowires, allow *S. oneidensis* cells to directly transfer electrons to denitrifying bacteria [16]; and 2) mediated electron transfer, in which soluble electron shuttles, such as flavins, serve as intermediaries that enable electron transfer between *S. oneidensis* and other denitrifying bacteria [18]. However, these transfer mechanisms are unlikely to fully represent the role that *S. oneidensis* plays in natural environments, considering its abundance of *S. oneidensis* and the unrestricted diffusion of electron shuttles. In denitrifying communities, *S. oneidensis* is rarely dominant, for example, it represented 2.51% of the 16S rRNA abundance detected in a denitrifying community that had been artificially inoculated with *S. oneidensis* [18]. Given that interspecies electron transfer occurs when abundant electron-donating bacteria typically exceeds 10% abundance [19], a significant enhancement of denitrification with a low abundance of *S. oneidensis* may imply that another mechanism also contributes to boosting the denitrifying activity.

Here, we demonstrate that membrane vesicles (MV) produced by *S. oneidensis* MR-1 enhanced the denitrification growth of various denitrifying bacteria. MV, which are nanoparticles ranging from 20–400 nm in size, are produced by a wide variety of bacteria, contain various biomolecules, and are used to transport nutrients, nucleotides, and proteins among cells [20]. MV are currently considered as mediators of interspecies interactions that are maintained in the environment for extended periods in complex polymicrobial communities [21–24] and are also found in denitrifying communities [25]. Further, the MV of *S. oneidensis* are components of nanowires, which are proposed as key mediators that connect bacteria and promote denitrification [11, 16, 17] because of their redox-active properties [26]. Proteomic analysis showed that these MV contain a significant amount of proteins related to respiration and cytochrome activity, similar to that found in the outer membrane of *S. oneidensis* cells [27]. Although MV produced from a cytochrome-abundant mutant have been shown to enhance biofilm formation and increase conductivity within biofilms [28], the impact that *S. oneidensis* MV have on other functions and interspecies interactions remains unclear.

In this study, we examined the impact that *S. oneidensis* MV have on the denitrification growth of five bacterial species and investigated their interactions, especially with *Pseudomonas aeruginosa* PAO1. We conducted a transcriptome analysis, tracked denitrification metabolites using isotopes, and performed experiments with loss-of-function mutants to elucidate the process by which MV enhance denitrification growth. Moreover, we discussed the attachment process of *S. oneidensis* MV with denitrifying bacteria by comparing the physical parameters of MV–bacteria interactions. Overall, our study findings indicate that *S. oneidensis* MV can play a notable role in polymicrobial communities, where they serve as enhancers of denitrifying bacteria in a species-specific manner.

## Materials and Methods

### Bacteria and growth

We used *Shewanella oneidensis* MR-1, *Pseudomonas aeruginosa* PAO1, *Pseudomonas stutzeri* NBRC14165, *Paracoccus aminovorans* NBRC16711, *Paracoccus denitrificans* PD1222 and *Achromobacter xylosoxidans* NBRC15126. The *S. oneidensis* mutant strains, Δ*fccA*Δ*cctA* and LS527, were used in the previous study [14, 29]. These strains, except *S. oneidensis*, were cultured on Luria–Bertani (LB) agar plates and in LB liquid medium at 37 °C. *S. oneidensis* wild-type and mutant strains were cultured on LB agar plates and in LB liquid medium at 30 °C.

To record the growth curves of these strains in the denitrification condition, we measured the optical density (OD) at 660 nm using an automated OD meter (TVS062CA, Advantec, Japan). Cells at the exponential phase were inoculated into fresh LB medium with and without 100 mM KNO_3_ at an initial OD of 0.1 in L-type test tubes. The test tubes were sealed with butyl rubber stoppers to culture in anoxic conditions, and the headspaces were replaced with helium gas (purity >99.999%). The test tubes were incubated at 37°C with vertical rotation at 30 rpm. OD was recorded every 5 minutes and the maximum growth speeds were calculated from five consecutive data points coinciding with the most rapid period of growth for each individual culture.

### Isolation and quantification of membrane vesicles

We isolated the MV produced by *S. oneidensis* and its mutants following previously described methods [30]. Briefly, *S. oneidensis* cells were cultured until they reached an OD of 0.3. Mitomycin C was then added at a concentration of 100 ng mL^-1^, mimicking the MV biogenesis route of *S. oneidensis* triggered by *P. aeruginosa* [31]. The culture was centrifuged and filtered through a 0.22 µm filter to separate the MV from the cells. The filtered supernatant was then ultracentrifuged at 150,000× g for 1 hour at 4 °C using Type 45 Ti rotor (Beckman Coulter) to collect the MV. The MV were resuspended in PBS solution. The structure of the MV samples was confirmed (Supplementary Figure S1). The amount of MV was compared by FM 1-43 staining (Thermo Fisher Scientific, Waltham, MA). The fluorescence of the sample with FM 1-43 was measured by a microplate reader (BioTek SYNERGY H1 microplate reader, Winooski, VT) with an excitation wavelength of 472 nm and an emission wavelength at 580 nm.

The MV produced by *P. aeruginosa* were prepared as described previously [32]. Briefly, *P. aeruginosa* cells were cultured until they reached an OD of 0.3. Mitomycin C was added at a concentration of 100 ng mL^-1^ and incubated for another 6 hours. The culture was centrifuged and filtered through a 0.45 µm filter. The supernatant was then ultracentrifuged at 150,000× g for 1 hour at 4 °C using Type 45 Ti rotor (Beckman Coulter), and then resuspended in PBS solution.

### Microbial adhesion to hydrocarbon test

The relative hydrophobicity of cell surface was assessed using the microbial adhesion to hydrocarbon (MATH) test [33, 34]. Briefly, bacterial cells were harvested during the exponential phase and washed three times with saline. Cell suspensions were prepared by adjusting the OD to 0.7 using saline (referred to as OD_before_), yielding a total volume of 3 mL for each sample. Subsequently, 1 mL of xylene was added to the cell suspension in a glass test tube. The suspension and oil were vortexed for 2 minutes at approximately 2700 rpm using a Vortex-Genie 2 (Scientific Industries, USA). The test tube was then left undisturbed for 10 minutes to allow the oil phase to separate and rise to the surface. After separation, the aqueous phase was carefully extracted using a pipette, ensuring minimal disturbance to the hydrocarbon layer. The optical density of the aqueous phase was measured (referred to as OD_after_). The MATH score (%) was calculated to quantify the relative cell adhesion to the hydrocarbon phase (formula 1)

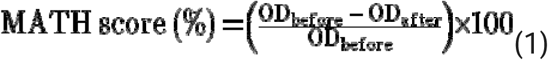

### Zeta potential

The zeta potential of the bacterial cell surface was measured following established methods [35]. Briefly, bacterial cells were harvested during the exponential growth phase and washed twice with saline. The cell suspensions were prepared in a 50 mM NaCl solution and diluted to 1/10 of the initial cell concentration. This was further diluted by 100-fold using Milli-Q water. The diluted suspensions were then filled into electrode cells. Zeta potential measurements were performed using a nanoparticle analyzer (HORIBA nanoPartica SZ-100V2, Kyoto, Japan) under electrode potentials of 3 to 4 V.

### Microscopic observation

We observed the structure of *S. oneidensis* MV and their interaction with *P. aeruginosa* cells using a transmission electron microscope (TEM) H7650 (Hitachi High-Technologies Corporation, Tokyo, Japan). To observe the interaction of MV with bacteria, *P. aeruginosa* cells were incubated with *S. oneidensis* MV for 250 minutes and then washed twice with PBS. The samples were diluted and placed on TEM grids (Cu400CN, ALLIANCE Biosystems, Osaka, Japan). The samples were washed with Milli-Q water and then stained using an EM stainer (Nissin EM, Japan).

To observe the interaction of MV-bacteria in fluorescence microscopy, we used non-stained *P. aeruginosa* cells and FM 1-43 stained MV. *S. oneidensis* MV were stained with FM 1-43 for 1 hour and then washed twice with PBS. Non-stained *P. aeruginosa* cells were incubated with FM 1-43 stained MV for 250 minutes and then washed twice with PBS. Both bright field and fluorescence microscopy images of these samples were taken using a fluorescence microscope equipped with a 100x/1.30 EC Plan-Neofluar objective lens (Carl Zeiss, Jena, Germany). The fluorescence microscopy images were observed under a condition with a green filter (excitation wavelength: 488 - 512 nm, emission wavelength: 520 - 550 nm).

### Transcriptome analysis

We investigated the transcriptome of *P. aeruginosa* cultured with or without *S. oneidensis* MV following the method described in the previous study [36, 37]. Briefly, cells were collected at the exponential phase, and RNA was isolated in the hot-phenol extraction, and purified with DNaseI and RNeasy Mini Kit (Quagen). RNA samples were sent to an external facility (Macrogen) for library preparation using the TruSeq Stranded Total RNA Library Preparation Kit (Illumina) and the Ribo-Zero Plus Microbiome rRNA Depletion Kit (Illumina). Sequencing was performed using the NovaSeq X Plus (Illumina). Transcriptome data analysis was performed as described in a previous study [36, 37]) with minor modifications. *P. aeruginosa* PAO1 reference genome data was downloaded from NCBI RefSeq database (accession number: NC_002516.2) and used for this analysis. Low-quality read removal and adapter trimming of the raw FASTQ data were performed using fastp software (version 0.23.4) [38] with “-q 30 -- detect_adapter_for_pe” options. Prior to mapping, an index file of the *P. aeruginosa* PAO1 genome FASTA file was generated using the genome’s GFF file and STAR (version 2.7.11a) [39] with “--runMode genomeGenerate –sjdbGTFfeatureExon gene – genomeSAindexNbases 10” options. The cleaned FASTQ data were then mapped to the genome using STAR with “--outSAMtype BAM SortedByCoordinate --alignIntronMax 1 --outFilterMultimapNmax 50” options. The resulting BAM files were indexed using the index function of Samtools (version 1.18) [40] with default options. The number of mapped reads to each gene were counted using featureCounts (version 2.0.6) [41] with “-p –countReadPairs -O -M -C -t gene -g locus_tag -s 2” options. Differentially expressed gene analysis was conducted using edgeR (version 4.0.16) (version 3.32.1). Low-count genes were filtered using the filterByExpr function with default options. Normalization was performed using the normLibSizes function with the trimmed mean of M-values method. Common dispersion, trended dispersions and tagwise dispersions were estimated with estimateDisp function. A quasi-likelihood F-test was performed using the glmQLFit functions to identify differentially expressed genes. q-values were adjusted using the topTags function with Benjamini and Hochberg method. Sequence data obtained in this study were deposited in the DNA Data Bank of Japan (DDBJ) Sequence Read Archive (DRA) under accession numbers PRJDB19804.

### Quantification of nitrogen oxides and nitrogen gas dynamics with acetate

To assess the denitrification process of *P. aeruginosa* with and without *S. oneidensis* MV, we used a minimal medium containing 1 mM K^15^NO_3_ and 2 mM acetate as described elsewhere [42]. *P. aeruginosa* cells cultured in LB with 100 mM KNO_3_ with and without *S. oneidensis* MV under anoxic conditions until exponential phase were harvested and washed twice with the minimal medium. The cells were then inoculated into the minimal medium at an initial OD of 1.0 and incubated at 37°C. We measured nitrate, nitrite, nitric oxide, nitrous oxide, and nitrogen gas following the methods described in the previous study [43]. Briefly, nitrate and nitrite were measured using colorimetric methods: the brucine-sulfanilic acid method for nitrate and the sulfanilamide-naphthyl ethylenediamine method for nitrite [44]. Acetate was measured using HPLC (Shimadzu, Kyoto, Japan) equipped with the Shim-pack SCR-102H column (Shimadzu, Kyoto, Japan). ^15^NO, ^15^N_2_O, and ^15^N_2_ were measured using a GCMS-QP 2010 SE gas chromatograph-mass spectrometer (Shimadzu, Kyoto, Japan).

In these conditions, *P. aeruginosa* cells utilize acetate and nitrate as an electron donor and an electron acceptor, respectively, where 8 electrons are produced by the oxidation of one molecule of acetate (formula 2) and 10 electrons are consumed by the reduction of two molecules of nitrate to one molecule of nitrogen gas during the denitrification process (formula 3).

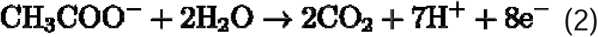

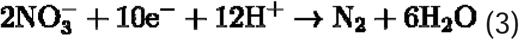

From these reactions, we estimate the ratio of electrons consumed by denitrification, based on acetate consumption (n_acetate_) and nitrogen gas production (n_N2_) (formula 4).

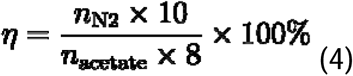

### Quantitative and statistical analyses

We performed all statistical tests in the python environment (python 3.9) with the SciPy library (https://scipy.org/). For pairwise comparisons, we assessed statistical significance between means using two-sample two-sided welch tests. When appropriate, we corrected p-values for multiple comparisons using the Bonferroni-Holm method. We reported the statistical test and the sample size (n) for each test in the results section, where all sample sizes are the number of independent biological replicates.

## Results

### Membrane vesicles produced by *Shewanella oneidensis* enhance bacterial denitrifying growth

*S. oneidensis* MR-1 cells have been shown to improve the denitrification performance of denitrifying bacteria when grown in co-culture [16–18]. To determine whether this improvement affects the growth of *P. aeruginosa*, we measured its growth in co-culture with *S. oneidensis* under both oxic and anoxic conditions (Supplementary Figure S2). Under oxic conditions, the growth rate in co-culture was almost identical to that in monoculture. While, consistent with previously reported improvements in denitrification [16–18], the growth rate under anoxic conditions was significantly higher in co-culture compared with that in monoculture (p_anoxic_ = 6.7 × 10^-5^; two-sample two-sided Welch test for μ_max_ [n ≥5]). Because the growth rate in co-culture should not exceed the growth rate in monoculture when two species utilize nutrient sources without specific interactions [45], the observed growth enhancement would be due to the presence of factors in the co-culture that supplement bacterial growth under anoxic conditions.

A plausible explanation for this enhancement involves interactions mediated by MV in the co-culture. MV production is induced by small compounds secreted from *P. aeruginosa* [31, 46], and *S. oneidensis* MV are likely involved in improving denitrification. To investigate the effect that *S. oneidensis* MV have on *P. aeruginosa* growth, we measured its nitrate-dependent growth after supplementation with *S. oneidensis* MV (Figure 1A). The growth of *P. aeruginosa* supplemented with *S. oneidensis* MV was significantly faster than that of the controls without MV (μ_max − MV_: 0.32 ± 0.05, μ_max + 4xMV_: 0.46 ± 0.03) (p = 6.0 × 10^−7^; two-sample two-sided Welch test for μ_max_ [n ≥10]). The maximum cell density (OD_max_) was also higher in the presence of MV (OD_max−MV_: 1.29 ± 0.11, OD_max + MV_: 1.71 ± 0.13) (p = 3.1 × 10^-7^; two-sample two-sided Welch test for OD_max_ [n ≥10]). These effects were confirmed at all concentrations of MV tested (single ∼ quadruple concentrations of MV isolated from *S. oneidensis* culture), indicating nitrate-dependent growth was enhanced by MV at physiological concentration level. Consistent with the enhancement of denitrification growth, NO_3_ ^−^ was more completely consumed (57.6 ± 11.2 mM consumption with MV supplementation; 23.8 ± 8.6 mM consumption without MV supplementation for 10 h of cultivation) (p = 3.9 × 10^−2^; two-sample two-sided Welch test for the maximum consumption rate [n = 3]) (Figure 1B). Even though *S. oneidensis* can grow in anoxic conditions with nitrate as an electron acceptor, MV produced by *P. aeruginosa* did not significantly affect the growth rate of *S. oneidensis* (Supplementary Figure S3). These results indicate that *S. oneidensis* MV specifically enhanced the nitrate-dependent growth of *P. aeruginosa*, likely through roles distinct from those of *P. aeruginosa* MV.

**Figure 1.**
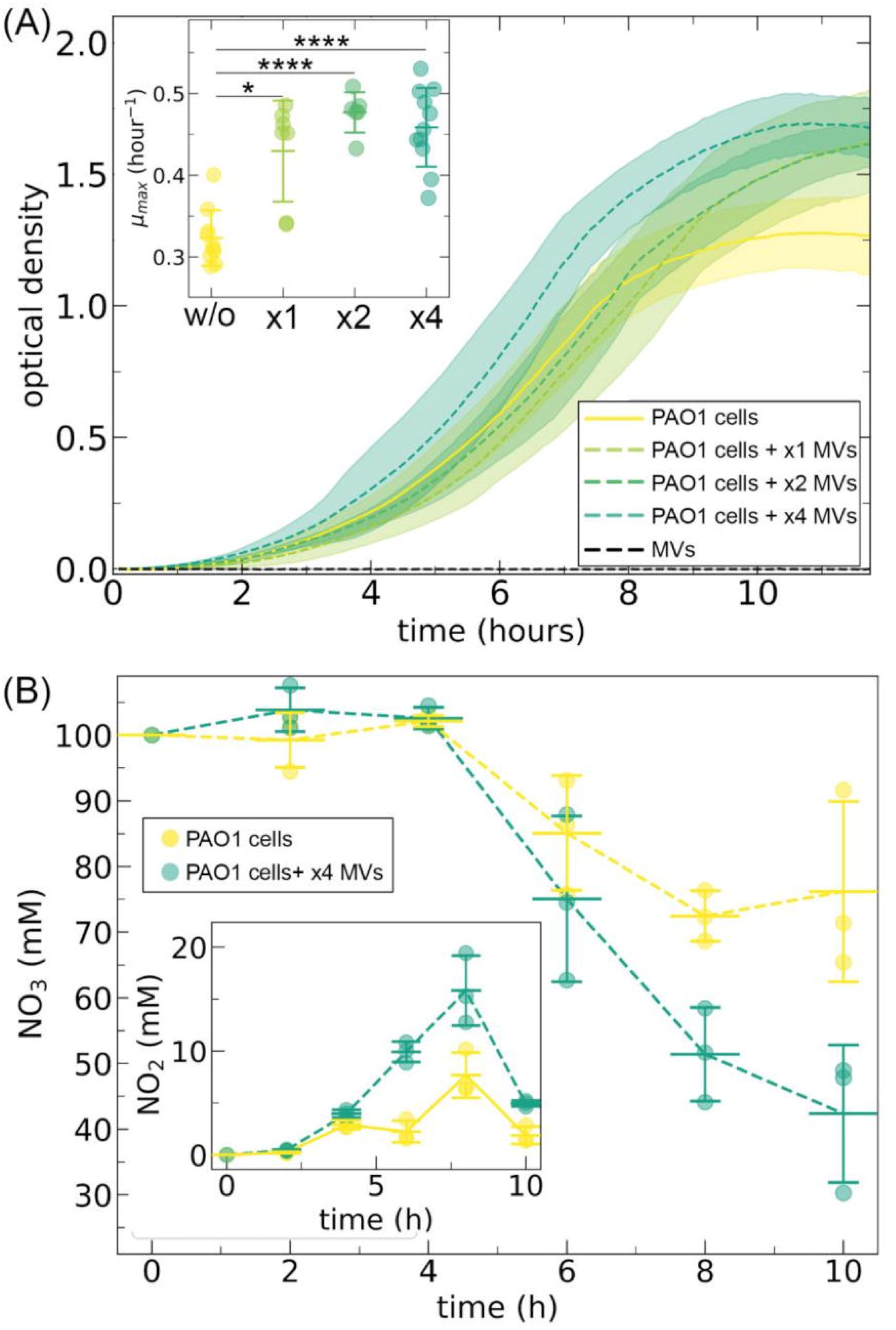
Membrane vesicles produced by *Shewanella oneidensis* enhance the denitrification growth of *Pseudomonas aeruginosa* PAO1. (A) *Pseudomonas aeruginosa* growth curve of cultures grown without (w/o; yellow) and with membrane vesicles (MV) produced by *Shewanella oneidensis* (yellow-green, dashed line) under anoxic conditions. The *P. aeruginosa* cells were cultured with different concentrations of *S. oneidensis* MV in Luria–Bertani (LB) medium with 100 mM nitrate at 37°C under anoxic conditions. The ×1, ×2, and ×4 MV labels correspond to concentrations from the same volume of *S. oneidensis* LB cultures. Lines are averages of independent biological replicates (n ≥ 5) and the shaded regions are ± one standard deviation. The inset shows the maximum growth speeds (µ_max_), which were calculated from five consecutive data points corresponding to the most rapid growth period for each individual culture. (B) Nitrate consumption as a function of time for *P. aeruginosa* cultured w/o MV (yellow) and with ×4 MV (yellow-green, dashed line) in LB medium containing 100 mM nitrate. The inset shows nitrite residues in the medium. Each data point represents an independent experimental replicate (n = 3). The middle horizontal line indicates the mean, and the upper and lower horizontal lines represent ± one standard deviation.

We further investigated the effects that *S. oneidensis* MV have on other denitrifying bacterial species. Specifically, we examined the nitrate-dependent growth of other four species: *P. stutzeri*, *P. aminovorans*, *P. denitrificans*, and *A. xylosoxidans* (Figure 2 and Supplementary Figure S4). *S. oneidensis* MV significantly enhanced the nitrate-dependent growth of *P. stutzeri* and *P. aminovorans* cultures (p*_P. stutzeri_* = 9.2 × 10^−3^, p*_P. aminovorans_* = 4.0 × 10^−3^; two-sample two-sided Welch tests for μ_max_ [n ≥7]). Specifically, the maximum growth rates and cell densities were higher in the presence of *S. oneidensis* MV than in controls without MV supplementation (Figure 2A and 2B). In contrast, the nitrate-dependent growth of *P. denitrificans* and *A. xylosoxidans* did not significantly differ with or without *S. oneidensis* MV supplementation (p*_P. denitrificans_* = 0.87, p*_A. xylosoxidans_* = 0.56; two-sample two-sided Welch test for μ_max_ [n ≥7]). These results indicate that the enhancement of nitrate-dependent respiration caused by *S. oneidensis* MV is species-specific. When MR-1 MV were added under oxic conditions in the absence of NO_3_ ^−^, neither the maximum growth rate nor the cell density significantly increased for any of the strains; rather, MV exposure inhibited the aerobic growth of *P. aeruginosa* (μ_max_: 0.75 ± 0.03 with MV; 0.89 ± 0.05 without MV) (p = 5.2 × 10^−6^, two-sample two-sided Welch test for μ_max_ [n ≥8]) (Supplementary Figure S5). Collectively, these results indicate that MV can specifically affect nitrate-dependent respiration without acting as a nutrient source for respiration.

**Figure 2.**
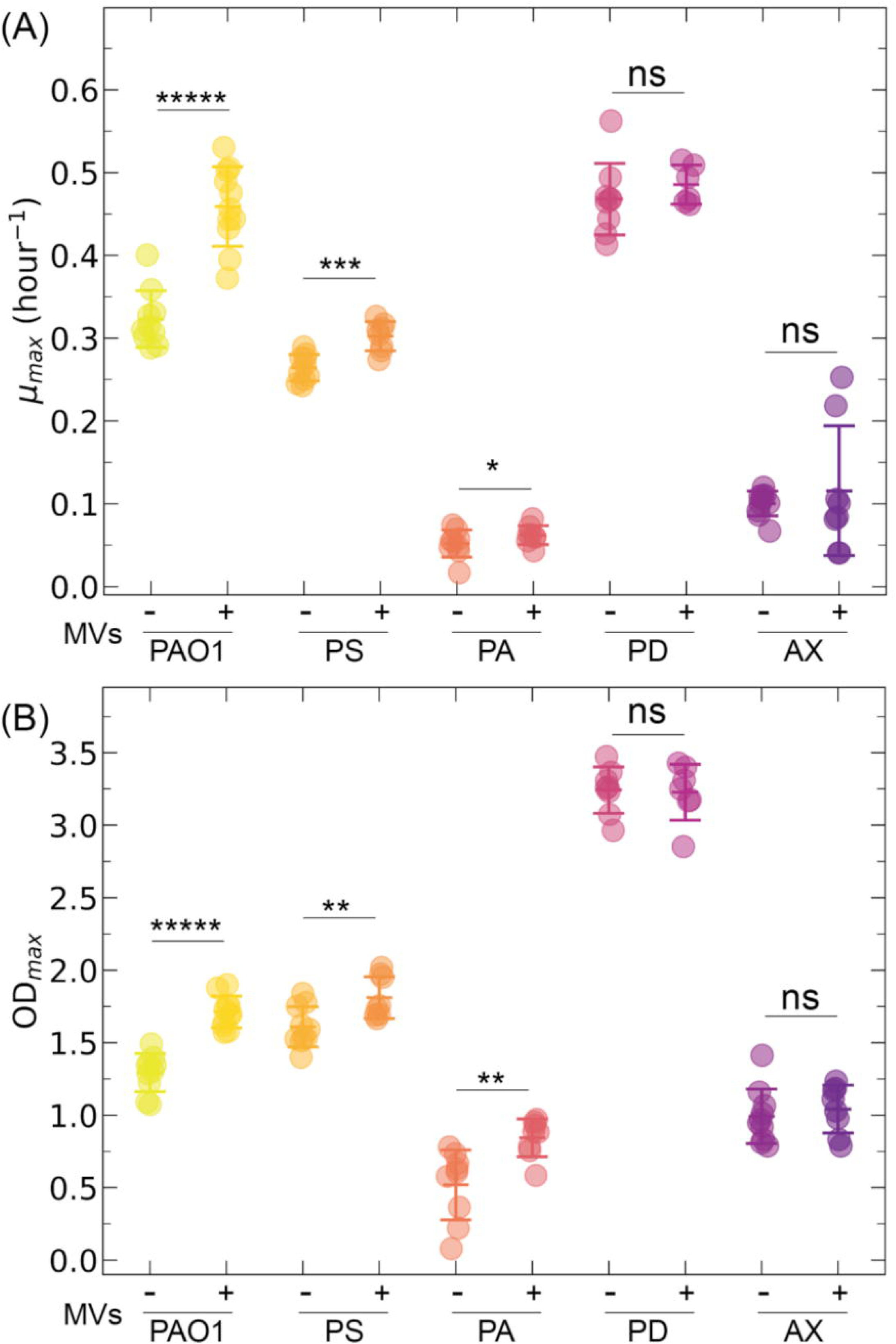
Membrane vesicles produced by *Shewanella oneidensis* enhance the denitrification growth of denitrifying bacteria. (A) Maximum growth speed (µ_max_) and (B) maximum cell density (OD_max_; OD = optical density) of *Pseudomonas aeruginosa* (PAO1), *Pseudomonas stutzeri* (PS), *Paracoccus aminovorans* (PA), *Paracoccus denitrificans* (PD), and *Achromobacter xylosoxidans* (AX) cultured without and with Membrane vesicles produced by *Shewanella oneidensis* (MV; ×4 MV concentrations) under anoxic conditions. Each data point represents an independent experimental replicate (n ≥5). The middle horizontal line indicates the mean, and the upper and lower horizontal lines represent ± one standard deviation. Asterisks indicate statistically significant differences in the means of the labelled groups, as calculated using two-sample two-sided Welch’s *t*-tests (ns, no significant difference; *, p <5.0 × 10^−2^; **, p <1.0 × 10^−2^; ***, p <1.0 × 10^−3^; ****, p <1.0 × 10^−4^; *****, p <1.0 × 10^−5^).

### Attachment between membrane vesicles and denitrifying bacteria

To elucidate the key factors dictating the species specificity of denitrification-dependent growth enhancement, we focused on the attachment of MV to bacterial surfaces. Although molecular mechanisms underlying the MV–bacteria interaction remain elusive [20], these interactions are likely influenced by the physical properties of membrane surfaces, specifically electrostatic interactions [47, 48]. Given that the surfaces of bacteria and MV are negatively charged, bacteria with surfaces with charges closer to neutrality should promote the attachment process [48, 49]. To examine whether electrostatic energy contributes to species specificity, we measured the zeta potentials of each denitrifying species used in this study (Supplementary Figure S6). The zeta potentials of these denitrifying bacteria were in the range of −40∼−80 mV, which are significantly negative values compared with those previously reported [49]. According to Tashiro et al., bacteria with zeta potentials less than −10 mV tend to create interface energy barriers, and potentials more than those with a zeta potential of −5 mV, which tend to create MV–bacteria interaction, matching the interaction energy calculation based on the Derjaguin–Landau–Verwey–Overbeek theory [49]. This suggests that under the tested conditions, electrostatic interactions between cells and MV do not contribute to the species specificity observed during nitrate-dependent growth enhancement.

Hydrophobicity is another key parameter that promotes stable interactions between phospholipid membranes [50]. To examine whether hydrophobicity contributes to species specificity, we conducted a MATH test that represents hydrophobic surface properties and compared the scores with the extent of growth enhancement observed. Owing to the technical difficulty associated with measuring MV, we measured the MATH score (MS) of *S. oneidensis* cells instead, resulting in a score of 49.7 ± 15.1, which suggests that *S. oneidensis* MV have hydrophobic surfaces (Figure 3A). Among the denitrifying bacteria, the strains that exhibited enhanced nitrate-dependent growth in response to MV (*P. aeruginosa* and *P. stutzeri*) showed high MATH scores (MS_PAO1_ = 50.9 ± 11.8, MS_PS_ =84.0 ± 4.1), whereas the other strains (*P. aminovorans*, *P. denitrificans*, and *A. xylosoxidans*) tended to show low MATH scores(MS_PA_ = 13.9 ± 9.6, MS_PD_ = 20.2 ±13.1, MS_AX_ = 1.2 ± 4.5). These results suggest that the species specificity occurs during nitrate-dependent growth enhancement arises from the hydrophobic attachment process.

**Figure 3.**
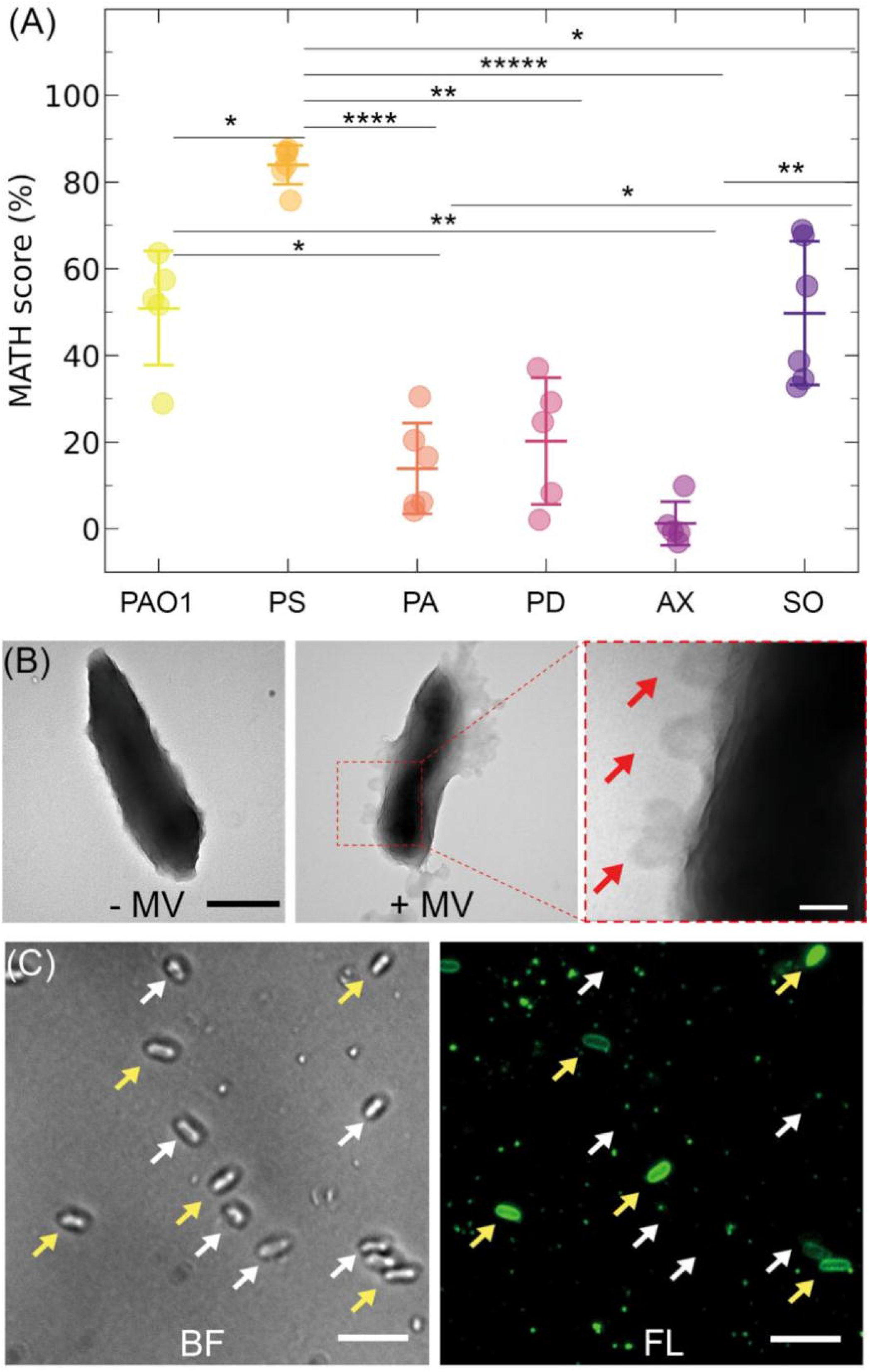
Elucidating the attachment process of membrane vesicles to denitrifying bacteria. (A) Microbial adhesion to hydrocarbon (MATH) scores (%) of *Pseudomonas aeruginosa* (PAO1), *Pseudomonas stutzeri* (PS), *Paracoccus aminovorans* (PA), *Paracoccus denitrificans* (PD), *Achromobacter xylosoxidans* (AX), and *Shewanella oneidensis* (SO). Cells were dispersed in saline solution before being vortexed with xylene. Each data point represents an independent experimental replicate (n = 5). The middle horizontal line indicates the mean, and the upper and lower horizontal lines represent ± one standard deviation. Asterisks indicate statistically significant differences in the means of the labelled groups, as calculated using two-sample two-sided Welch’s *t*-tests corrected with the Bonferroni–Holm method (*, p <5.0 × 10^−2^; **, p <1.0 × 10^−2^; ***, p <1.0 × 10^−3^; ****, p <1.0 × 10^−4^; *****, p <1.0 × 10^−5^). (B) Representative micrographs of *P. aeruginosa* cultured with *S. oneidensis* membrane vesicles (MV) viewed under a transmission electron microscopy. The expanded view is shown in the left panel. The white arrows highlight circular structures surrounding the *P. aeruginosa* cell, indicating the presence of *S. oneidensis* MV. Scale bars = 1 µm (left, black) and 100 nm (right, white). (C) Representative micrographs of *P. aeruginosa* cells cultured with *S. oneidensis* MV captured using bright field (left) and fluorescence (right) microscopy. The cells at high and low fluorescence are shown with yellow and white arrows, respectively. Scale bar = 5 µm.

To directly confirm the attachment process between MV and denitrifying bacteria, we observed that *S. oneidensis* MV were substantially attached to *P. aeruginosa* cells using TEM and fluorescence microscopy. In the TEM images, we identified spherical particles resembling MV surrounding *P. aeruginosa* cells in the samples incubated with *S. oneidensis* MV (Figure 3B). These spherical structures were absent in *P. aeruginosa* cells without *S. oneidensis* MV supplementation, indicating the association between *S. oneidensis* MV and *P. aeruginosa* cells. Multiple MV were observed to associate with the cell surface, rather than a single MV attaching to a single cell. Furthermore, fluorescence microscopy was performed to confirm the attachment process of *S. oneidensis* MV to *P. aeruginosa* cells. Before incubation of *S. oneidensis* MV and *P. aeruginosa* cells, the MV were stained with FM 1-43, a fluorescent dye for membranes. This result showed that the *P. aeruginosa* cells exhibited green fluorescence, indicating the presence of *S. oneidensis* MV associated with these cells (Figure 3C). Some *P. aeruginosa* cells exhibited high levels of green fluorescence, whereas others remained nonfluorescent, which is a characteristic of MV–bacteria interactions [24]. Taken together, these results indicate that *S. oneidensis* MV have an affinity for *P. aeruginosa* cells, supporting that MV directly interact with *P. aeruginosa* cells.

### Membrane vesicles produced by *Shewanella oneidensis* enhance denitrification reaction efficiency independently of transcriptional regulation

To investigate the mechanisms underlying the enhancement of denitrification that occurs after MV attachment, we measured the gene expression levels of *P. aeruginosa* with and without *S. oneidensis* MV supplementation using RNA-seq (Figure 4 and Supplementary Table 1). Our analysis revealed that 19 genes were upregulated (log_2_ fold change >1 and −log_10_ q-value >2) and 67 downregulated (log_2_ fold change <−1 and −log_10_ q-value >2) in the presence of *S. oneidensis* MV. Surprisingly, genes involved in denitrification, such as those encoding nitrogen oxide reductases and transporters, did not show significant differences in their expression levels with and without *S. oneidensis* MV supplementation. These results suggest that *S. oneidensis* MV enhance denitrification by influencing denitrifying activities without affecting their transcriptional regulation.

**Figure 4.**
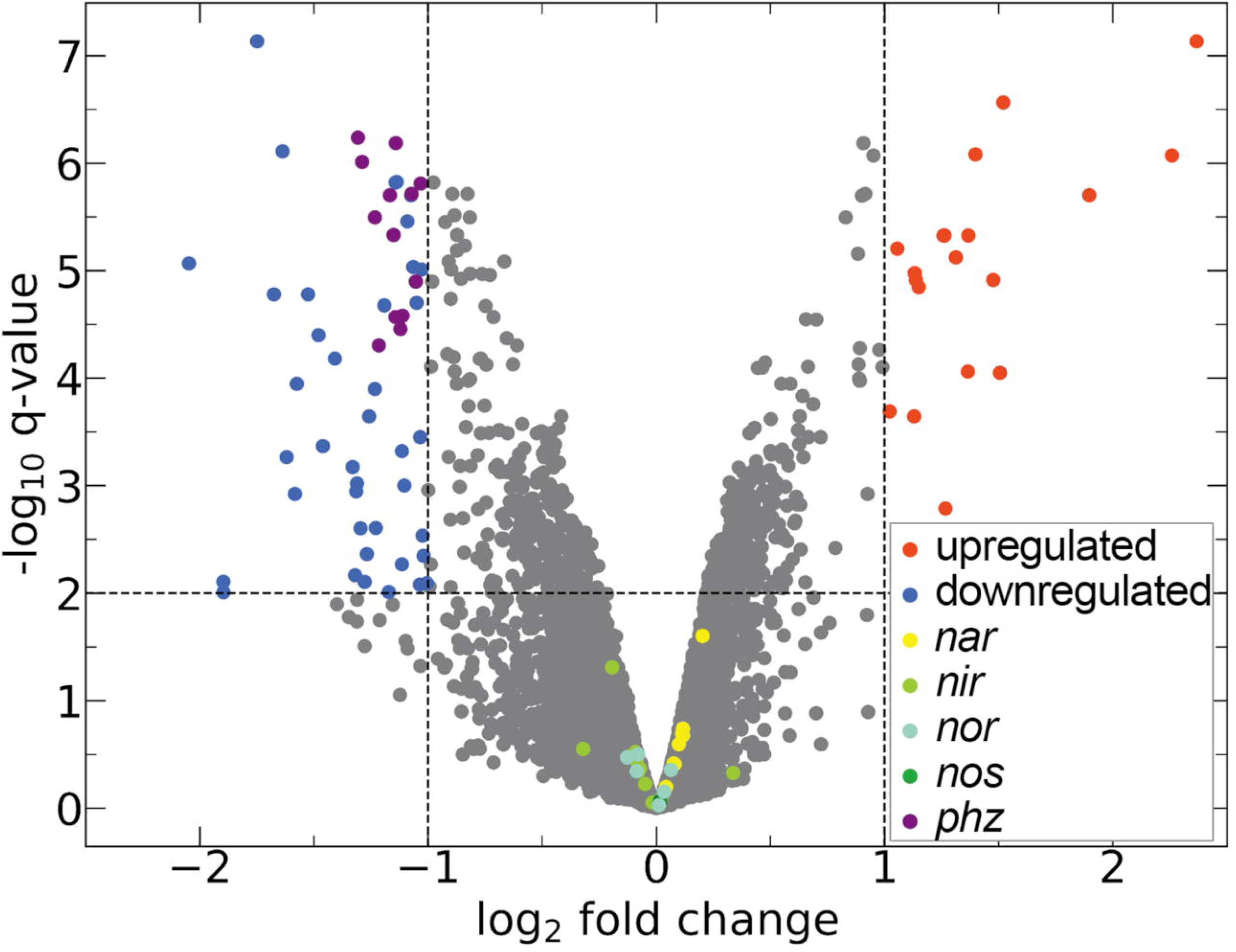
Denitrification gene cluster expression levels are not significantly different between cultures grown with and without membrane vesicles produced by *Shewanella oneidensis*. (A) Volcano plots indicating the differential gene expression levels of *Pseudomonas aeruginosa* cultured with Membrane vesicles produced *by Shewanella oneidensis* when compared with *P. aeruginosa* cultured (MV) without *S. oneidensis* MV. RNA was extracted from *P. aeruginosa* cells at mid-log phase, cultured both with and without *S. oneidensis* MV, for this analysis. The horizontal line represents a threshold where the −log_10_ q-value >2. The vertical lines represent thresholds for the log_2_ fold change, with values >1 or <−1. Genes that are significantly upregulated (log_2_ fold change >1 and −log_10_ q-value >2) are highlighted in red, whereas those significantly downregulated (log_2_ fold change <−1 and −log_10_ q-value >2) are highlighted in blue. Notably, the denitrification gene clusters (*nar*, *nir*, *nor*, and *nos*) and phenazine genes are shown in yellow, yellow-green, green, aquamarine, and purple, respectively.

In total, 42% (8 out of 19) of the upregulated genes were related to tRNA synthesis, indicating an increase in protein translation activity. Contrastingly, the downregulated genes were more diverse. In particular, expression levels of the *ccb_3_* cytochrome oxidase (*ccoN4*, PA4133, and PA4134) and phenazine synthesis gene clusters were significantly downregulated in *P. aeruginosa* cultures supplemented with MV. The *ccb_3_*cytochrome oxidase and phenazines are crucial for respiratory electron transport in *P. aeruginosa* cells, with cytochrome oxidase reducing oxygen to water and phenazines extracellularly shuttling electrons to maintain redox balance [51, 52].

To verify that *S. oneidensis* MV enhance nitrate-dependent growth independently of transcriptional regulation, we quantified denitrification metabolites under conditions where growth was minimized, leading to the estimation of denitrifying activities. We measured the concentrations of acetate, NO_3_^−^, NO_2_^−^, NO, N_2_O, and N_2_ using 1 mM isotopically labelled NO_3_^−^ (^15^NO_3_^−^) and 2 mM acetate as the sole carbon source under anoxic conditions in a minimal medium at an initial OD of 1.0 (Figure 5). The consumption rate of NO_3_^−^ was higher in *P. aeruginosa* cultures supplemented with MV (p = 6.2 × 10^−5^; two-sample two-sided Welch’s *t-*test [n = 5]), whereas NO_3_^−^ was completely consumed under both conditions. Consistent with NO_3_^−^ consumption, N_2_ was produced under both conditions, but its production rate was higher in the presence of MV (p = 6.8 × 10^−7^; two-sample two-sided Welch test [n = 5]) (Figure 5A). We detected denitrification intermediates, such as NO_2_^−^ and N_2_O, under both conditions; however, these concentrations were significantly lower compared with that of N_2_ (Supplementary Figure S7). Additionally, the concentration of acetate in *P. aeruginosa* cultures supplemented with *S. oneidensis* MV remained at approximately 0.4 mM at the end of the culture period, whereas it was completely consumed in *P. aeruginosa* cultures without *S. oneidensis* MV supplementation (Figure 5B). By estimating the denitrification efficiency normalized by acetate consumption, we found that it was significantly higher in the presence of *S. oneidensis* MV (p = 1.8 × 10^−4^; two-sample two-sided Welch test [n = 5]) (Figure 5B). These results indicate that *S. oneidensis* MV influence denitrifying activities, rather than the transcription of denitrification-related genes.

**Figure 5.**
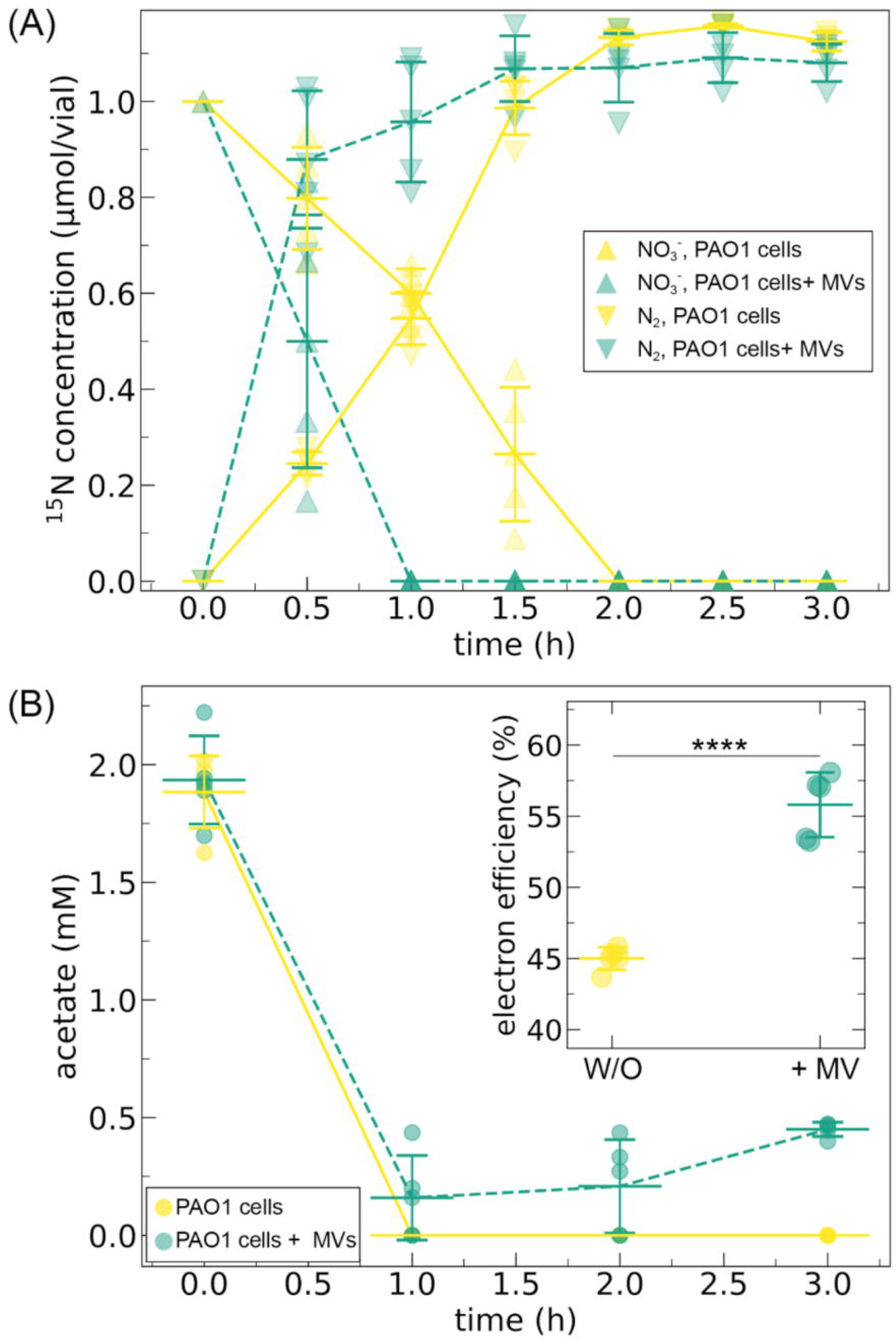
Membrane vesicles produced by *Shewanella oneidensis* enhance denitrification by *Pseudomonas aeruginosa* under conditions where the cells scarcely grew. (A) Dynamics of NO_3_^−^ reduction (solid line, upper triangle) and N_2_ production (dashed line, lower triangle) in *Pseudomonas aeruginosa* cultured without (w/o; yellow) and with Membrane vesicles produced by *Shewanella oneidensis* (MV; ×4 MV concentrations, green). (B) Acetate consumption in *P. aeruginosa* cultured w/o (yellow) and with *S. oneidensis* MV (green). The inset shows the ratio of electrons consumed by denitrification, which were estimated from N_2_ production and acetate consumption at 3 h (calculation formula described in Methods). The *P. aeruginosa* cells were inoculated at an optical density of 1.0 in minimal medium containing 1 mM ^15^KNO_3_ and 2 mM acetate at 37°C under anoxic conditions. Each data point represents an independent experimental replicate (n = 5). The middle horizontal line indicates the mean, and the upper and lower horizontal lines represent ± one standard deviation.

### *c*-Type cytochromes in membrane vesicles contribute to the enhancement of denitrification growth

MV can serve as cargo for transporting small molecules, including quorum-sensing signals and redox-active molecules [53, 54]. Soluble redox molecules are known to enhance denitrification activity [18, 55]. We hypothesized that MV containing electron shuttle-like flavins might similarly influence denitrification growth. Electron-shuttling flavins can be generated by *S. oneidensis*, including flavin mononucleotide (FMN), flavin adenine dinucleotide (FAD), and riboflavin (RF) [56]. To test this hypothesis, we examined the effects that these flavins have on the denitrification growth of *P. aeruginosa*. Contrary to our expectations, these molecules did not enhance denitrification growth under our experimental conditions; rather, they inhibited denitrification growth (p_FMN_ = 3.5 × 10^−7^, p_FAD_ = 2.4 × 10^−3^, p_RF_ = 1.3 × 10^−3^; two-sample two-side Welch tests for μ_max_ [n ≥5]), (Figure 6A and Supplementary Figure S8). To investigate whether this inhibitory effect occurs across various redox-active compounds with differing potentials, we examined artificial redox molecules with physiologically relevant redox potentials, including anthraquinone-2-sulfonate (+30 mV vs. Standard Hydrogen Electrode, SHE), 2-hydroxy-1,4-naphthoquinone (−150 mV vs. SHE), and methylene blue (−11 mV vs. SHE). Additionally, we examined phenazine-1-carboxylic acid and pyocyanin, which are typical redox-active secondary metabolites of *P. aeruginosa*. Similar to that observed for flavins, these compounds inhibited denitrification growth (Figure 6A and Supplementary Figure S8). These results suggest that electron shuttles, such as flavins, do not contribute to enhancing denitrification growth through MV.

**Figure 6.**
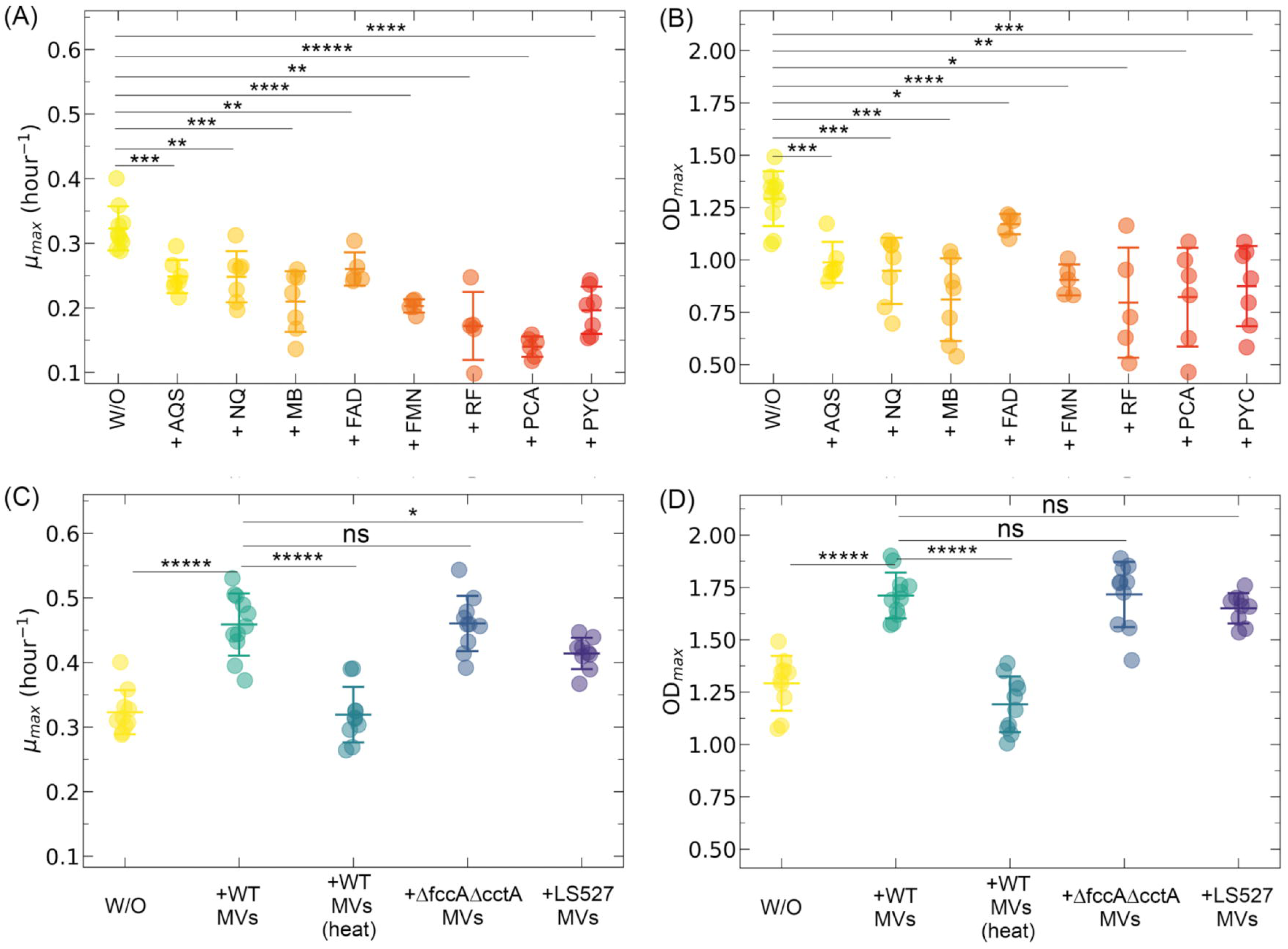
Impact of redox-active molecules on the enhancement of denitrification. (A) Maximum growth speed (µ_max_) and maximum reaching points (OD_max_; OD = optical density) of *Pseudomonas aeruginosa* growth in the presence of 100 µM electron mediators. AQS: anthraquinone-2-sulfonate; NQ: 2-hydroxy-1,4-naphthoquinone; MB: methylene blue; FAD: flavin mononucleotide; FMN: flavin adenine dinucleotide; RF: riboflavin; PCA: phenazine-1-carboxylic acid; PYC: pyocyanin. (C) The µ_max_ and (D) OD_max_ of *P. aeruginosa* growth in the presence of membrane vesicles (MV; ×4 MV concentrations) from *Shewanella oneidensis* mutants lacking cytochrome-related genes (Δ*fccA*Δ*cctA*, LS527) and inactive wild-type (WT) MV, which were treated at 80°C for 3 h. Each data point represents an independent experimental replicate (n >5). The middle horizontal line indicates the mean, and the upper and lower horizontal lines represent ± one standard deviation. Asterisks indicate statistically significant differences in the means of the labelled groups, as calculated using two-sample two-sided Welch’s *t*-tests (ns, no significant difference; *, p <5.0 × 10^−2^; **, p <1.0 × 10^−2^; ***, p <1.0 × 10^−3^; ****, p <1.0 × 10^−4^; *****, p <1.0 × 10^−5^).

MV produced by *S. oneidensis* contain redox active proteins, such as *c*-type cytochromes that exhibit redox activity [11, 27, 30]. To investigate the contribution of MV components to denitrification growth, we measured the nitrate-dependent growth using the heat-treated WT MV that were treated at 80°C for 3 h. The heat-treated MV failed to enhance growth compared to untreated WT MV (p_heat-treated MV_ = 1.1 × 10^−6^; two-sample two-sided Welch test for μ_max_ [n ≥9]). Since heat treatment denatures and inactivates MV components such as proteins, this suggests that MV components contribute to denitrification enhancement. To further investigate the contribution of cytochromes contained in MV, MV isolated from a mutant strain (LS527), which lacks outer-membrane *c*-type cytochromes [14], were added to *P. aeruginosa* cultures. Although MV produced by this mutant enhanced the growth rate and maximum cell density of *P. aeruginosa* compared with those measured under the MV-free condition, the maximum growth rate was slightly lower than that after supplementation with wild-type (WT) MV (p_LS527 MV_ = 1.6 × 10^−2^; two-sample two-sided Welch *t-*test for μ_max_ [n ≥9])(Figure 6B and Supplementary Figure S8). This indicates that *c*-type cytochromes are partially involved in enhancing denitrification. In contrast, MV sourced from the Δ*fccA*Δ*cctA* mutant, where the genes encoding *c*-type cytochromes in the periplasmic space were deleted, enhanced denitrifying growth of *P. aeruginosa* in an almost identical level with that of the WT (p_Δ*fcc*Δ*cctA* MV_ = 0.94; two-sample two-sides Welch *t-*test for μ_max_ [n ≥9]). Overall, these results demonstrate that MV component such as outer-membrane *c*-type cytochromes contribute to denitrification enhancement, likely beyond serving solely as electron mediators.

## Discussion

By analysing the denitrification-dependent growth of *P. aeruginosa* cultured with MV produced by *S. oneidensis* (Figure 1), we provide evidence that MV are involved in respiratory processes. The extracellular extensions of *S. oneidensis* are well-known to exhibit the functions of redox-active components [11, 12, 26, 27, 57]. Previous studies on denitrifying bacterial interaction have primarily focused on direct electron transfer via extracellular extensions [16–18] or mediated electron transfer via electron shuttles, such as flavins secreted from *S. oneidensis* [18]. Our results demonstrate that *S. oneidensis* MV enhanced denitrification growth without transporting electrons from cells. Using exogenous electron mediators and mutant MV, we demonstrated that *c*-type cytochromes in MV are one of the key components involved in denitrification enhancement (Figure 6). This effect may surpass the redox activities of conventional electron mediators. These findings imply a potential role for MV in modulating bacterial respiration, possibly by delivering specialized components that interact with recipient cells to promote respiratory/metabolic activity.

Our findings reveal that *S. oneidensis* MV enhanced denitrification growth in a species-specific manner (Figure 2). Unlike conventional approaches that utilize soluble redox molecules to promote denitrification, MV demonstrate interactions with specific microbial species, and under our experimental conditions, enhance denitrification more significantly. This species-specific enhancement of denitrification growth could confer a competitive advantage to certain species, enabling their dominance within denitrifying communities. In natural environments, MV are more abundant than bacterial cells and contribute to crucial roles in bacterial communities, including nutrient transport and signalling [53, 58]. Environments rich in denitrifying bacteria, such as lakes, soils, and wastewater treatment plants where *S. oneidensis* thrives, are likely reservoirs of diverse MV [25]. MV-based respiratory interactions may be a common form of bacterial communication in nature, with potential implications for biogeochemical cycles. Further research into the specific composition of *S. oneidensis* MV and the molecular mechanisms underlying MV–bacteria interactions could enable more precise applications of MV in biotechnological and environmental contexts.

Several studies have investigated interspecies interactions between cells and MV, indicating that physical properties, such as surface hydrophobicity and electrical charge, are key factors that influence the attachment process [49, 59–61]. However, it is unclear whether these parameters are related to the biological functions of MV. We found that bacteria with highly hydrophobic surfaces tended to be susceptible to denitrification enhancement (Figure 3A), suggesting that hydrophobicity is a key mediating factor. Thus, technologies that modulate the physical properties of MV or cell, such as those developed through synthetic biological and chemical approaches [62–64], could serve as simple and promising strategies to control bacterial respiration/metabolism with species specificity. These approaches do not require detailed molecular-level insights into MV–bacteria interactions; they only necessitate the development of methodologies to adjust bacteria or MV hydrophobicity. This would be advantageous when compared with designing novel chemicals with species specificity, which faces considerable challenges and requires molecular-level insights to design chemical structures suitable for target bacterial transporters and enzymes sensitive to water solubility.

MV–bacteria interactions are likely complex and may involve various steps, including molecular recognition with cell appendages, MV–bacteria fusion, uptake of MV components, etc. Thus, it is important to clarify the mechanisms underlying the denitrification enhancement that occurs after MV attach to bacteria. Our study demonstrated that *c*-type cytochromes in MV are key components that enhance denitrification. The *c*-type cytochromes are thought to be localized on the surface of *S. oneidensis* MV [30], suggesting that their role in denitrification may require close interactions between MV surfaces and the periplasmic space of *P. aeruginosa*. This interaction appears to influence the physiological responses of *P. aeruginosa*. Although the detailed mechanisms underlying denitrification enhancement after MV–bacteria adsorption are unclear, this predicts that the presence of *S. oneidensis* MV triggers specific regulatory mechanisms in *P. aeruginosa*, leading to alterations in gene expression. Gene expression analysis revealed that *P. aeruginosa* downregulates genes related to electron transport, such as the *ccb*_3_ cytochrome oxidase gene (*ccoN4*) and phenazine synthesis clusters, when exposed to *S. oneidensis* MV. Although *ccb*_3_ cytochrome oxidase primarily functions to transfer electrons to oxygen, its expression remains elevated under anoxic conditions, potentially to maintain redox balance in low-oxygen environments [65]. Previous studies have shown that deletion mutants lacking *ccb*_3_ cytochrome oxidase (Δ*ccoN1*Δ*ccoN2*) exhibit enhanced denitrification in *P. aeruginosa* [66]. Furthermore, we exogenously added phenazine to *P. aeruginosa* cultures, which led to the inhibition of denitrification growth. Consistent with these findings, *S. oneidensis* MV suppressed *ccoN4* expression and phenazine biosynthesis, enhancing denitrification. These findings suggest that MV modulate electron transfer, balancing redox states and easing respiratory competition. This highlights their ecological role in optimizing microbial metabolism and cooperation in polymicrobial communities.

In conclusion, we demonstrated that MV produced by *S. oneidensis* enhanced the denitrification growth of various denitrifying bacteria, clarifying their versatility and specificity in influencing respiratory processes across bacterial species. Given the ubiquity of MV in natural environments, this study underscores their potential role as key mediators of interspecies interactions in microbial communities. In ecosystems with a high abundance of denitrifying bacteria, such as freshwater sediments, soils, and wastewater treatment systems, MV-based respiratory interactions could actively shape microbial community dynamics and contribute to the maintenance of ecosystem functions. The ability of MV to specifically enhance denitrification may promote microbial cooperation and competition, ultimately influencing species composition and ecological stability. Furthermore, by elucidating the ecological significance of MV in microbial respiration, this study provides novel insights into their potential role in biogeochemical cycles. These findings not only enhance our understanding of microbial interactions but also offer practical applications for optimizing metabolic efficiency in biotechnological processes.

## Acknowledgements

We thank Dr. David R. Johnson for his valuable assistance with word expression and comments; Editage (www.editage.jp) for English language editing; Prof. Dr. Liang Shi and Prof. Dr. Johannes Gescher for kindly providing the LS527 and Δ*fccA*/Δ*cctA* mutants, respectively.

## Author contributions

K.T. and Y.T. designed the study. K.T., R.T., T.S., M.K., and Y.T. performed the experimental work and analyzed the data. T. Y. and O. M. contributed to the data analysis. K.T. and Y.T. wrote the first draft of the manuscript. All authors contributed to the editing and finalizing of the manuscript.

## Data availability

Analysis scripts and processed data have been deposited at figshare (10.6084/m9.figshare.28386062). Sequence data have been deposited in the DDBJ (accession numbers are shown in Materials and Methods)

