## Supplementary file for "Membrane vesicles of *Shewanella oneidensis* MR-1 enhance denitrification growth in a species-specific manner"

*Corresponding Author

**This file includes:**

Tables S1 and S2

Figures S1 - S8

### Supplementary Information

#### Supplementary Table S1. The list of upregulated genes (log_2_ fold change > 1 and -log_10_ q-value > 2) of *P. aeruginosa* in the presence of *S. oneidensis* MV compared with *P. aeruginosa* in the absence of *S. oneidensis* MV.

| Locus tag | Gene | Function | Log FC | FDR |
| --- | --- | --- | --- | --- |
| PA0433 | - | hypothetical protein | 1.06 | 6.2E-06 |
| PA0434 | - | hypothetical protein | 1.52 | 2.7E-07 |
| PA0435 | - | hypothetical protein | 1.40 | 8.3E-07 |
| PA0439 | - | dihydropyrimidine dehydrogenase | 1.90 | 2.0E-06 |
| PA0440 | - | oxidoreductase | 1.50 | 8.9E-05 |
| PA0441 | *dht* | D-hydantoinase/dihydropyrimidinase | 2.26 | 8.5E-07 |
| PA0443 | - | transporter | 1.48 | 1.2E-05 |
| PA0444 | - | allantoate amidohydrolase | 1.37 | 4.7E-06 |
| PA1796.3 | - | tRNA-Leu | 1.13 | 2.3E-04 |
| PA1796.4 | - | tRNA-His | 1.02 | 2.0E-04 |
| PA2097 | - | flavin-binding monooxygenase | 1.36 | 8.7E-05 |
| PA2098 | - | esterase | 1.27 | 1.6E-03 |
| PA2757 | - | hypothetical protein | 2.37 | 7.4E-08 |
| PA3094.1 | *-* | tRNA-Asp | 1.15 | 1.4E-05 |
| PA3094.2 | - | tRNA-Asp | 1.13 | 1.1E-05 |
| PA3094.3 | - | tRNA-Val | 1.14 | 1.2E-05 |
| PA4541.1 | - | tRNA-Lys | 1.31 | 7.5E-06 |
| PA4541.2 | - | tRNA-Pro | 1.26 | 4.7E-06 |
| PA4541.3 | - | tRNA-Asn | 1.26 | 4.7E-06 |

#### Supplementary Table S2. The list of downregulated genes (log_2_ fold change > 1 and -log_10_ q-value > 2) of *P. aeruginosa* with *S. oneidensis* MV compared to *P. aeruginosa* without *S. oneidensis* MV.

| Locus tag | gene | function | Log FC | FDR |
| --- | --- | --- | --- | --- |
| PA0736a | - | hypothetical protein | -1.11 | 4.7E-04 |
| PA0737 | - | hypothetical protein | -1.57 | 1.1E-04 |
| PA0843 | *plcR* | phospholipase C accessory protein PlcR | -1.17 | 9.7E-03 |
| PA1219 | *-* | hypothetical protein | -1.03 | 4.8E-02 |
| PA1874 | *-* | hypothetical protein | -1.14 | 1.5E-06 |
| PA1899 | *phzA2* | phenazine biosynthesis protein PhzA | -1.12 | 3.5E-05 |
| PA1900 | phzB2 | phenazine biosynthesis protein PhzB | -1.05 | 1.3E-05 |
| PA1901 | *phzC2* | phenazine biosynthesis protein PhzC | -1.03 | 1.5E-06 |
| PA1902 | *phzD2* | phenazine biosynthesis protein PhzD | -1.29 | 9.7E-07 |
| PA1903 | phzE2 | phenazine biosynthesis protein PhzE | -1.31 | 5.8E-07 |
| PA1904 | *phzF2* | trans-2,3-dihydro-3-hydroxyanthranilate isomerase | -1.14 | 6.5E-07 |
| PA1905 | *phzG2* | pyridoxamine 5'-phosphate oxidase | -1.11 | 2.6E-05 |
| PA2048 | - | hypothetical protein | -1.35 | 1.7E-02 |
| PA2066 | - | hypothetical protein | -1.03 | 3.5E-04 |
| PA2067 | - | hydrolase | -1.02 | 2.9E-03 |
| PA2068 | - | major facilitator superfamily transporter | -1.23 | 1.3E-04 |
| PA2069 | - | carbamoyl transferase | -1.53 | 1.7E-05 |
| PA2122 | - | hypothetical protein | -1.41 | 6.6E-05 |
| PA2123 | - | transcriptional regulator | -1.05 | 2.0E-05 |
| PA2167 | - | hypothetical protein | -1.09 | 3.3E-02 |
| PA2175 | - | hypothetical protein | -1.28 | 3.1E-02 |
| PA2176 | - | hypothetical protein | -1.21 | 1.8E-02 |
| PA2262 | - | 2-ketogluconate transporter | -1.01 | 8.0E-03 |
| PA2322 | - | gluconate permease | -1.03 | 9.7E-06 |
| PA2324 | - | hypothetical protein | -1.31 | 1.8E-02 |
| PA2375 | - | hypothetical protein | -1.32 | 6.8E-03 |
| PA2385 | *pvdQ* | acyl-homoserine lactone acylase PvdQ | -1.15 | 1.3E-02 |
| PA2397 | *pvdE* | pyoverdine biosynthesis protein PvdE | -1.04 | 8.3E-03 |
| PA2413 | *pvdH* | diaminobutyrate--2-oxoglutarate aminotransferase | -1.46 | 4.3E-04 |
| PA2512 | *antA* | anthranilate dioxygenase large subunit | -1.27 | 4.3E-03 |
| PA2513 | *antB* | anthranilate dioxygenase small subunit | -1.90 | 7.8E-03 |
| PA2514 | *antC* | anthranilate dioxygenase reductase | -1.12 | 8.8E-02 |
| PA2570 | *lecA* | PA-I galactophilic lectin | -1.58 | 1.2E-03 |
| PA2939 | - | aminopeptidase | -1.07 | 2.0E-06 |
| PA3195 | *gapA* | glyceraldehyde 3-phosphate dehydrogenase | -1.19 | 2.1E-05 |
| PA3221 | *csaA* | molecular chaperone CsaA | -1.31 | 9.5E-04 |
| PA3222 | - | hypothetical protein | -1.33 | 6.7E-04 |
| PA3444 | - | alkanesulfonate monooxygenase | -1.30 | 2.5E-03 |
| PA3445 | - | hypothetical protein | -2.05 | 8.6E-06 |
| PA3446 | - | NAD(P)H-dependent FMN reductase | -1.48 | 4.0E-05 |
| PA3447 | - | ABC transporter ATP-binding protein | -1.90 | 9.8E-03 |
| PA3448 | - | ABC transporter permease | -1.10 | 2.8E-02 |
| PA3449 | - | hypothetical protein | -1.31 | 1.1E-02 |
| PA3514 | - | ABC transporter ATP-binding protein | -1.02 | 4.5E-03 |
| PA3517 | - | adenylosuccinate lyase | -1.11 | 5.4E-03 |
| PA3518 | - | hypothetical protein | -1.26 | 2.3E-04 |
| PA3724 | *lasB* | elastase LasB | -1.06 | 9.2E-06 |
| PA3923 | - | hypothetical protein | -1.10 | 9.9E-04 |
| PA3937 | - | taurine ABC transporter ATP-binding protein | -1.32 | 1.1E-03 |
| PA3938 | - | taurine-binding protein | -1.68 | 1.7E-05 |
| PA4133 | - | cbb3-type cytochrome C oxidase subunit I | -1.64 | 7.8E-07 |
| PA4134 | - | hypothetical protein | -1.75 | 7.4E-08 |
| PA4137 | - | porin | -1.28 | 7.8E-03 |
| PA4141 | - | hypothetical protein | -1.14 | 1.5E-06 |
| PA4142 | - | secretion protein | -1.09 | 3.5E-06 |
| PA4159 | *fepB* | iron-enterobactin transporter periplasmic binding protein | -1.62 | 5.4E-04 |
| PA4209 | *phzM* | phenazine-specific methyltransferase | -1.22 | 5.0E-05 |
| PA4210 | *phzA1* | phenazine biosynthesis protein | -1.14 | 2.7E-05 |
| PA4211 | *phzB1* | phenazine biosynthesis protein | -1.23 | 3.2E-06 |
| PA4212 | *phzC1* | phenazine biosynthesis protein PhzC | -1.07 | 1.9E-06 |
| PA4213 | *phzD1* | phenazine biosynthesis protein PhzD | -1.29 | 9.7E-07 |
| PA4214 | *phzE1* | phenazine biosynthesis protein PhzE | -1.31 | 5.8E-07 |
| PA4215 | *phzF1* | trans-2,3-dihydro-3-hydroxyanthranilate isomerase | -1.14 | 6.5E-07 |
| PA4216 | *phzG1* | pyridoxamine 5'-phosphate oxidase | -1.15 | 4.7E-06 |
| PA4217 | *phzS* | hypothetical protein | -1.17 | 2.0E-06 |
| PA4306 | *flp* | type IVb pilin Flp | -1.23 | 2.5E-03 |
| PA4814 | *fadH2* | 2,4-dienoyl-CoA reductase | -1.40 | 1.3E-02 |

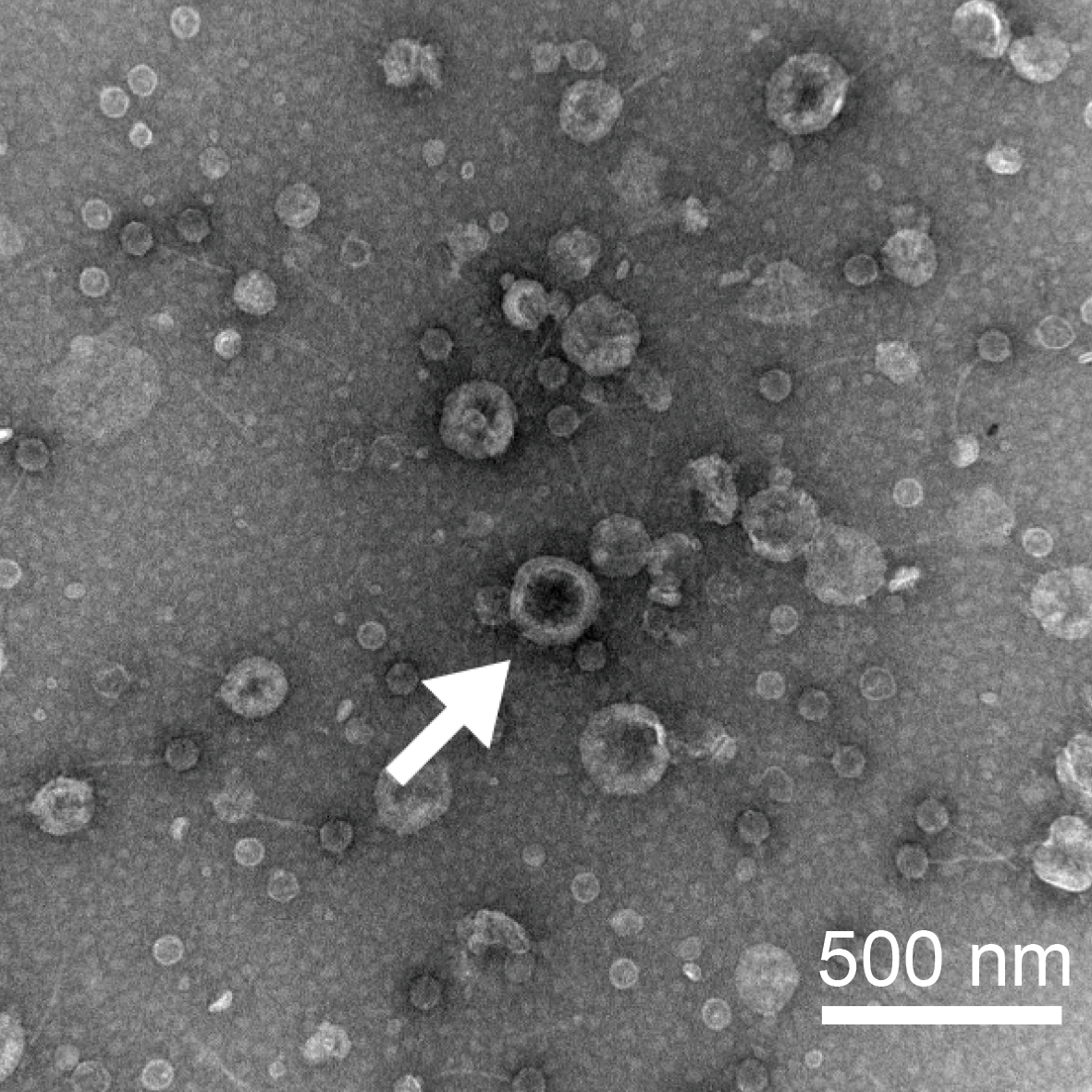

#### Supplementary Figure S1. A transmission electron microscopic image of membrane vesicles produced by *Shewanella oneidensis*

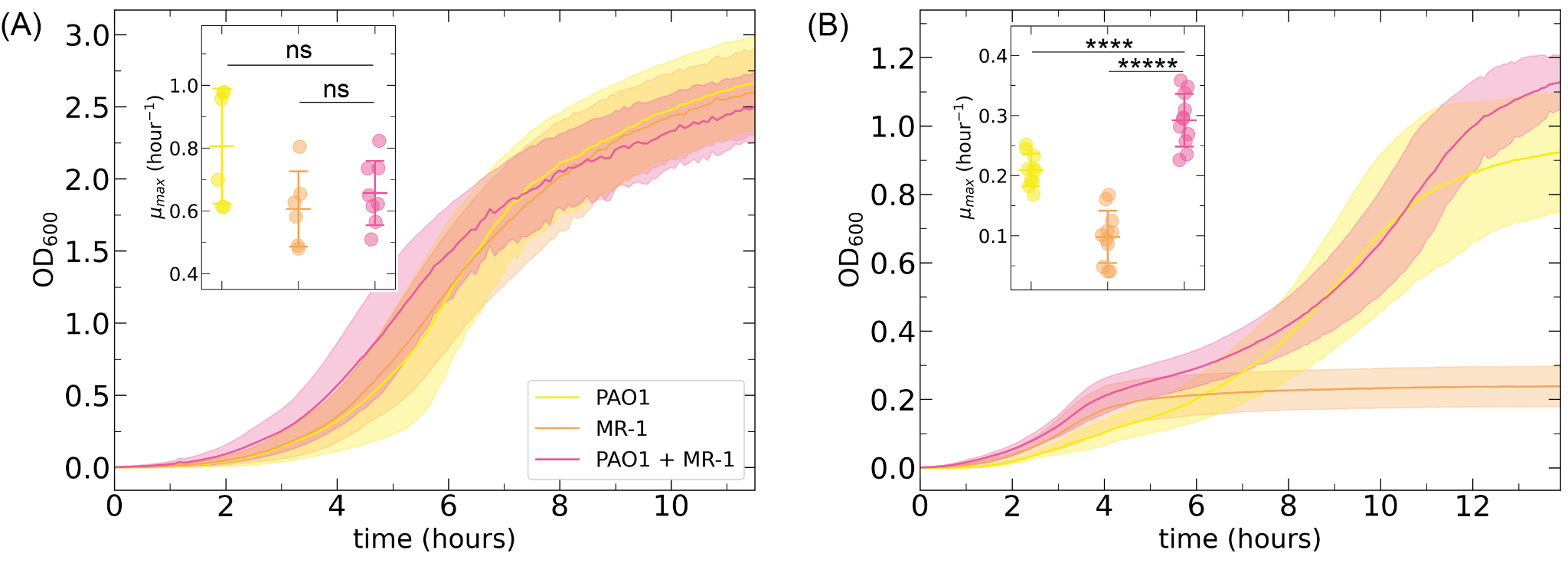

#### Supplementary Figure S2. Growth curve of mono- and co-culture for *Pseudomonas aeruginosa* and *Shewanella oneidensis* in oxic and anoxic condition

The growth curve of monoculture, *Pseudomonas aeruginosa* (yellow), *Shewanella oneidensis* (orange), and co-culture of *P. aeruginosa* and *S. oneidensis* (pink) in oxic (A) and anoxic (B) conditions. We culture them in LB medium at 30 ºC. Data points are averages of independent biological replicates (n ≥ 5) and the shaded regions are ±1 standard deviation. The inset shows the maximum growth speeds (µ_max_), which were calculated from five consecutive data points corresponding to the most rapid growth period for each culture. The middle horizontal line indicates the mean, and the upper and lower horizontal lines represent ±1 standard deviation. Asterisks indicate statistically significant differences in the means of the labeled groups calculated with two-sample two-sided welch test ns; no significant difference, **; p < 1.0 x 10-2, ***; p < 1.0 x 10-3, ****; p < 1.0 x 10-4, *****; p < 1.0 x 10-5)

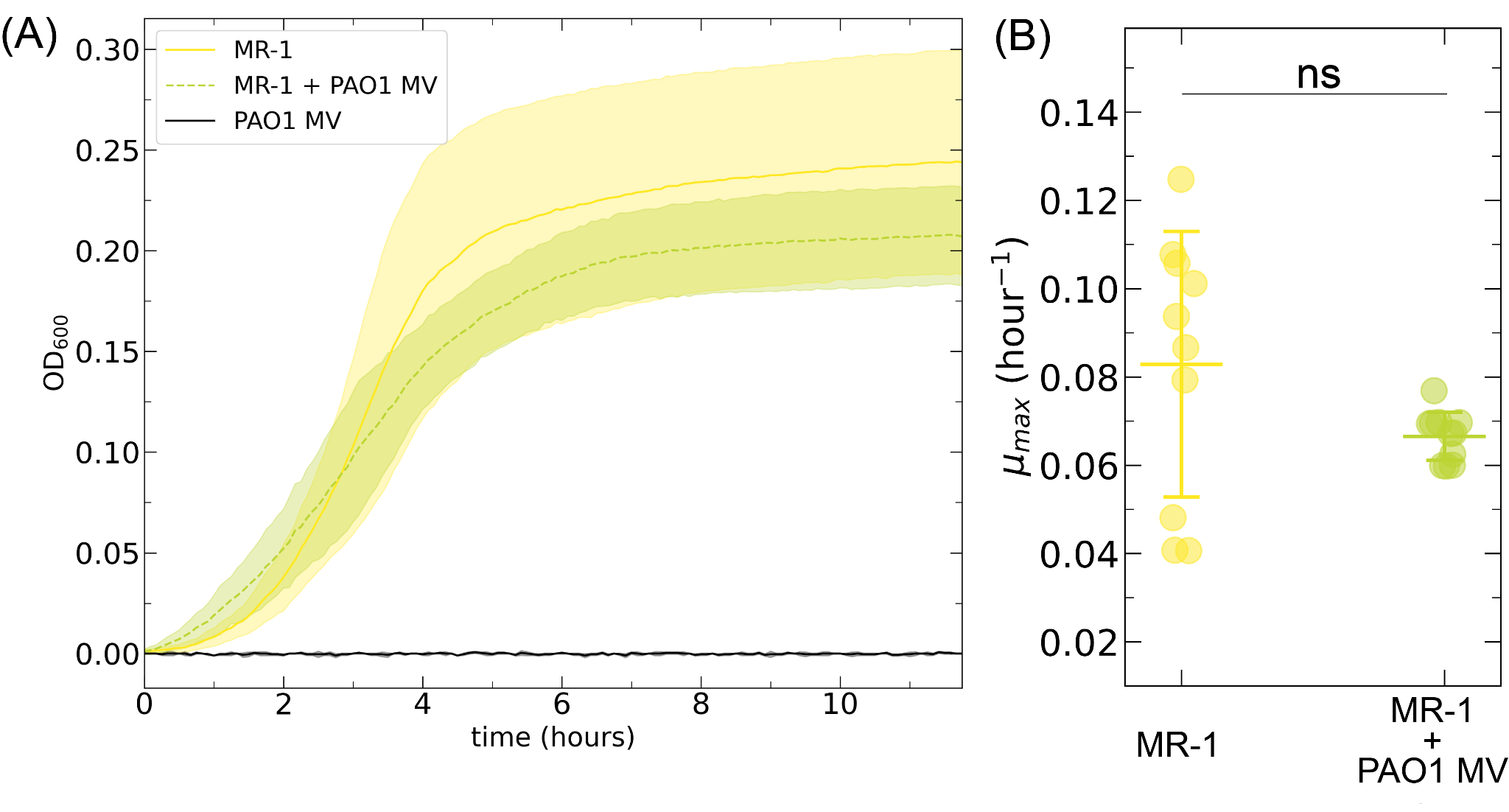

#### Supplementary Figure S3. *Shewanella oneidensis* growth with membrane vesicles produced by *Pseudomonas aeruginosa* under anoxic conditions

(A) *Shewanella oneidensis* growth curve without (yellow, solid line) and with membrane vesicle produced by *Pseudomonas aeruginosa* (green, dashed line) in the anoxic condition. We culture *S. oneidensis* cells with *P. aeruginosa* MV (x4 MV concentrations) in LB medium with 100 mM nitrate at 30 ºC under anoxic conditions. Data points are averages of independent biological replicates (n ≥ 5) and the shaded regions are ±1 standard deviation. (B) The maximum growth speeds (µ_max_), which were calculated from five consecutive data points corresponding to the most rapid growth period for each culture. The middle horizontal line is the mean and the upper and lower horizontal lines are ±1 standard deviation. The means of the labelled groups calculated with two-sample two-sided welch test are not significant differences.

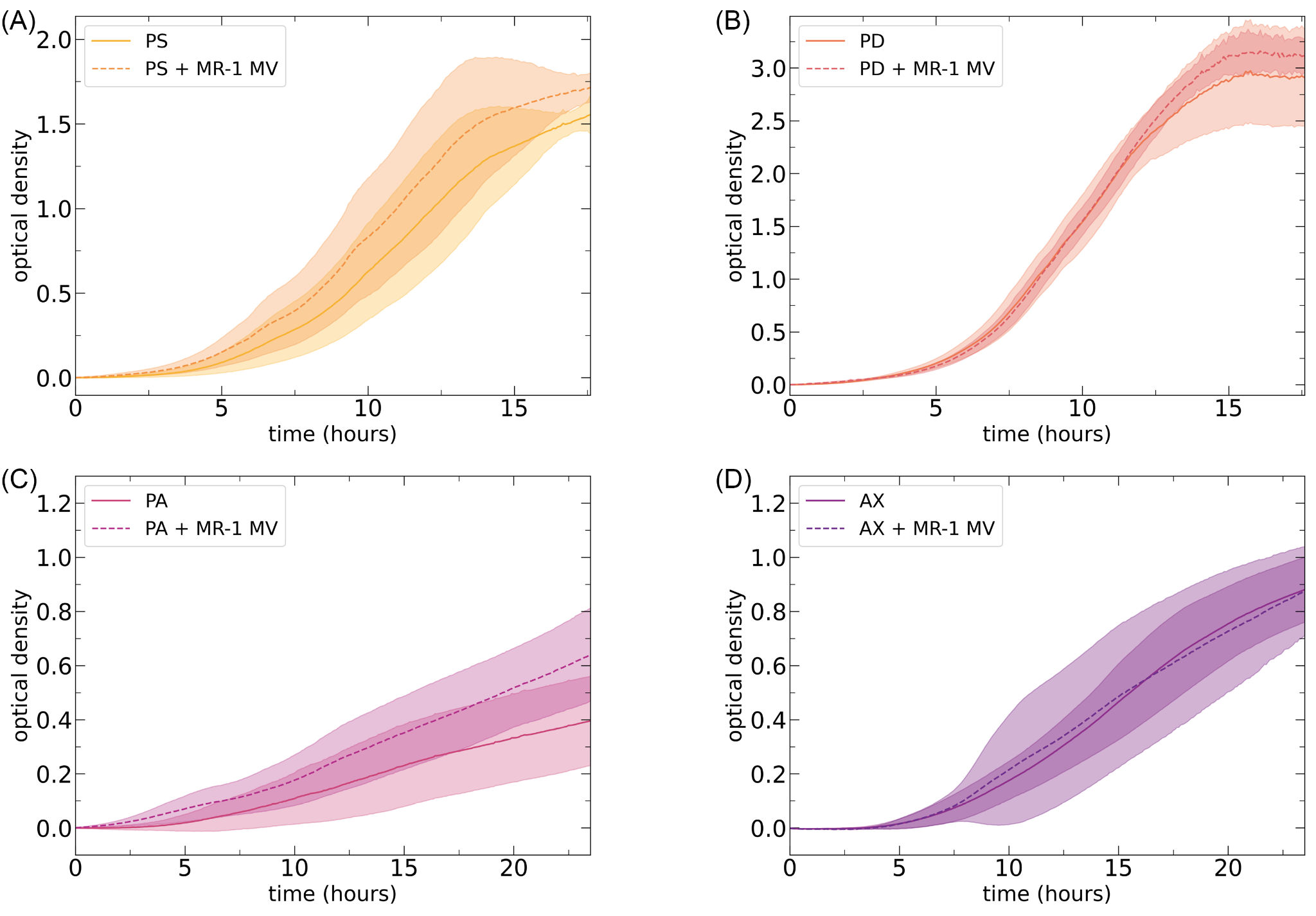

#### Supplementary Figure S4. Denitrifying bacteria growth with and without membrane vesicles produced by *Shewanella oneidensis* in anoxic condition

The growth curve of *Pseudomonas stutzeri* (PS), *Paracoccus aminovorans* (PA), *Paracoccus denitrificans* (PD), and *Achromobacter xylosoxidans* (AX) without (solid line) and with MV (dashed line). We cultured these species with *S. oneidensis* MV (x4 MV concentrations) in LB medium with 100 mM nitrate at 37 ºC in anoxic conditions. Data points are averages of independent biological replicates (n ≥ 5) and the shaded regions are ±1 standard deviation.

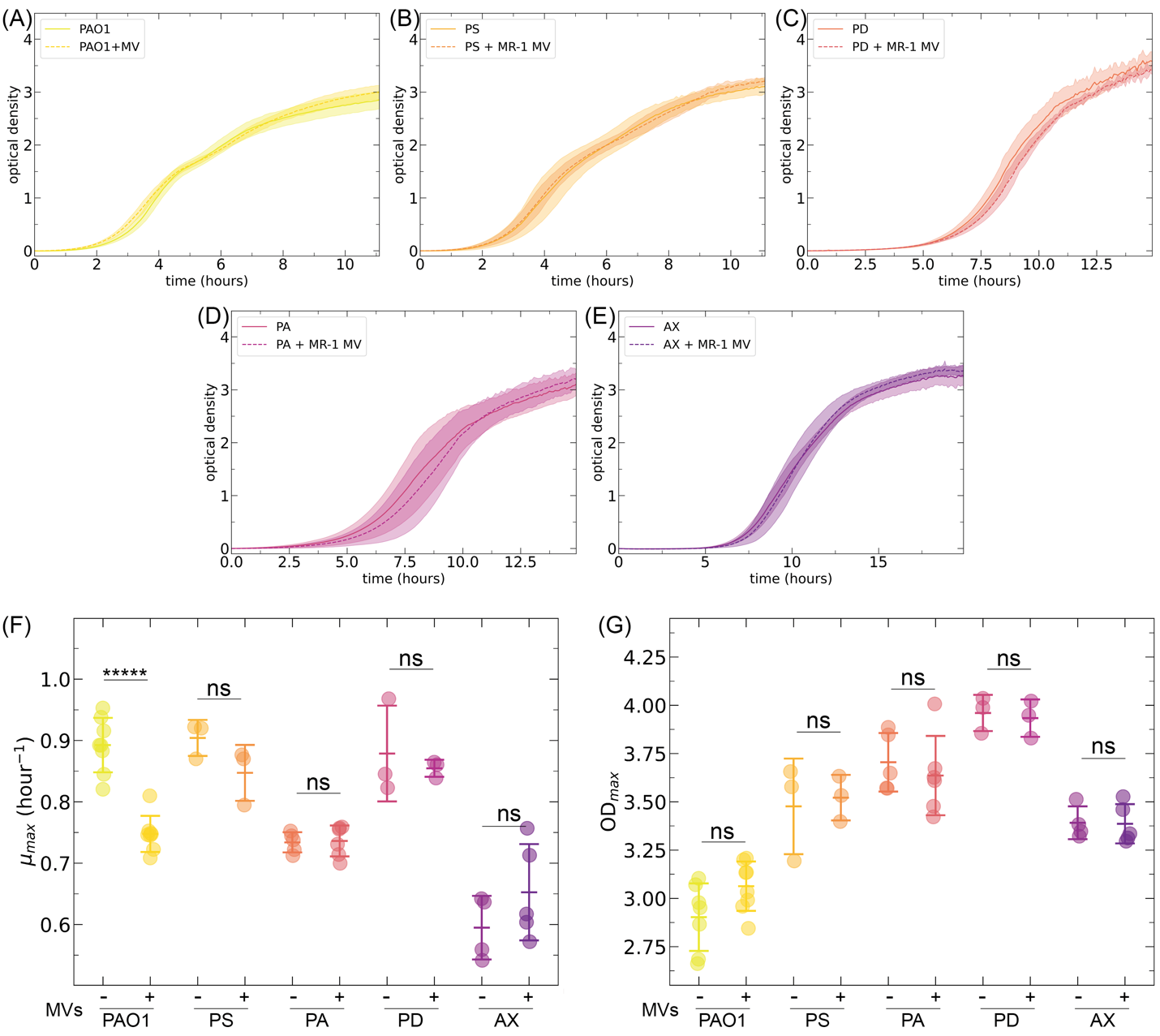

#### Supplementary Figure S5. Denitrifying bacteria growth with and without membrane vesicles produced by *Shewanella oneidensis* in oxic condition

(A-E) The growth curve of *Pseudomonas aeruginosa* (PAO1), *Pseudomonas stutzeri* (PS), *Paracoccus aminovorans* (PA), *Paracoccus denitrificans* (PD), and *Achromobacter xylosoxidans* (AX) without (solid line) and with MV (dashed line) in oxic condition. We culture these species with *S. oneidensis* MV (x4 MV concentrations) in LB medium at 37 ºC in oxic conditions. Data points are averages of independent biological replicates (n ≥ 5) and the shaded regions are ±1 standard deviation. (F) The maximum growth speed and (G) the maximum cell density (OD_max_) without and with *S. oneidensis* MV in oxic conditions. The maximum growth speeds were calculated from five consecutive data points coinciding with the most rapid period of growth for each culture. Each data point represents an independent experimental replicate (n ≥ 3). The middle horizontal line indicates the mean, and the upper and lower horizontal lines represent ±1 standard deviation. Asterisks indicate statistically significant differences in the means of the labelled groups calculated with two-sample two-sided welch test ns; no significant difference, *; p < 5.0 x 10^-2^, *****; p < 1.0 x 10^-5^).

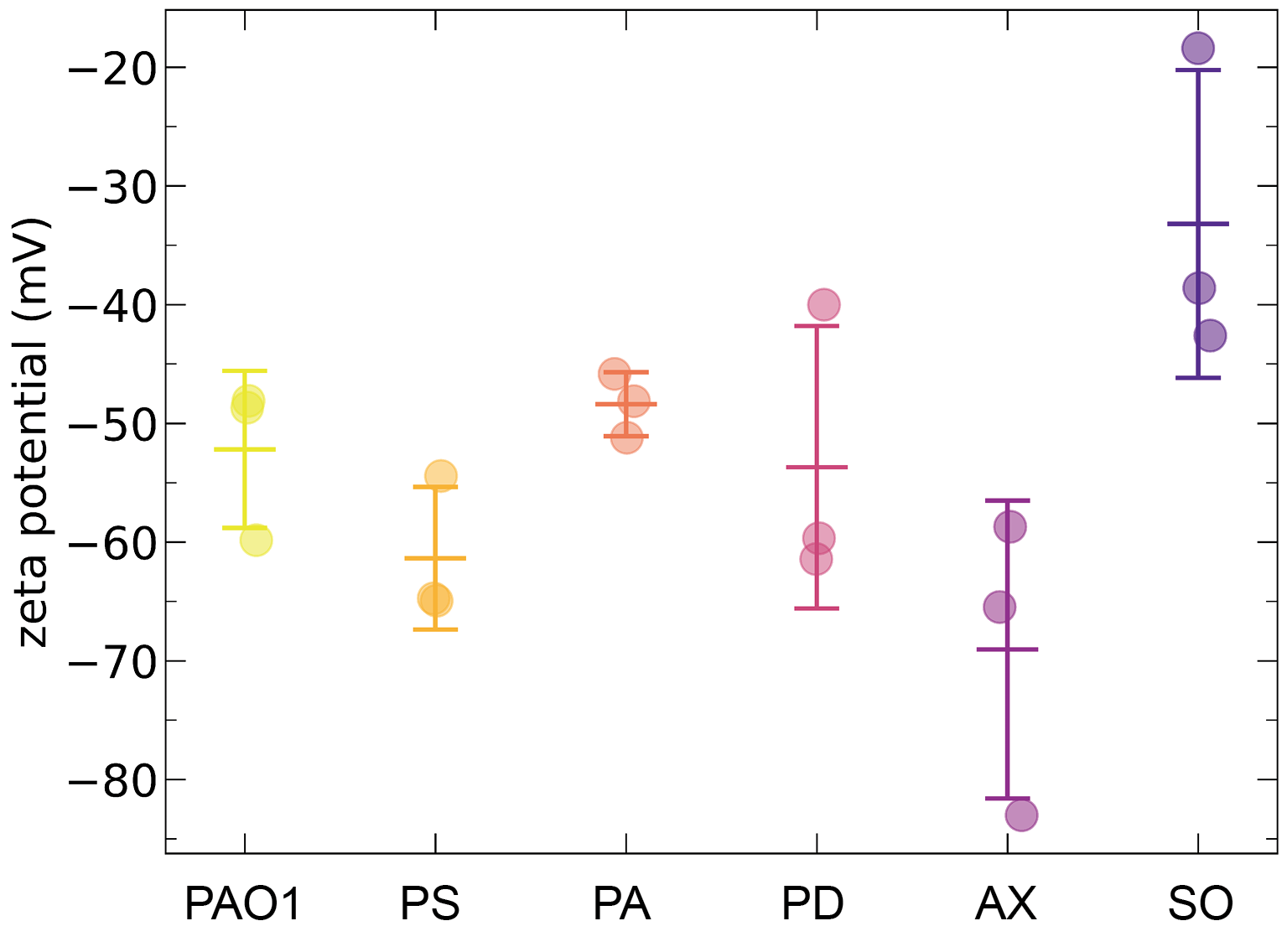

#### Supplementary Figure S6. Surface electrostatic properties of denitrifying bacteria and *Shewanella oneidensis*

The zeta potential of *Pseudomonas aeruginosa* (PAO1), *Pseudomonas stutzeri* (PS), *Paracoccus aminovorans* (PA), *Paracoccus denitrificans* (PD), *Achromobacter xylosoxidans* (AX), and *Shewanella oneidensis* (SO). Each data point represents an independent experimental replicate (n = 3). The middle horizontal line indicates the mean, and the upper and lower horizontal lines represent ±1 standard deviation.

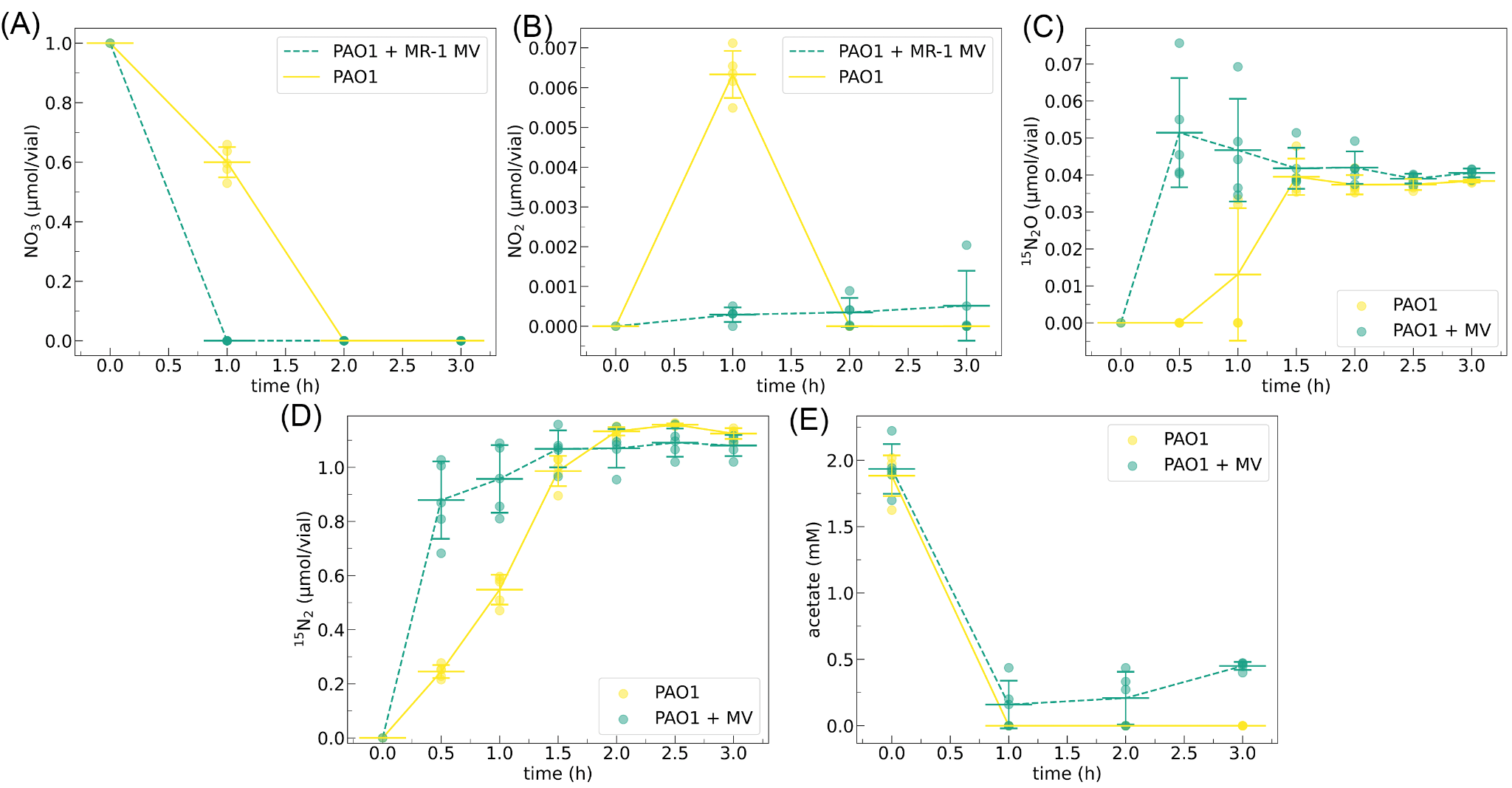

#### Supplementary Figure S7. Nitrogen compound profiles as a function of time in the minimal medium with acetate as the sole carbon source

Concentrations of substrates and products for *Pseudomonas aeruginosa* cultures without (yellow) and with (green) *S. oneidensis* MV (x4 MV concentrations). *P. aeruginosa* cells were cultured in the minimal medium containing 1 mM KNO_3_ and 2 mM acetate at 37°C under anoxic conditions. Data points are presented for (A) nitrate, (B) nitrite, (C) nitrous oxide, (D) nitrogen gas, and (E) acetate over time. Nitric oxide was not detected in GC-MS. Data points represent averages of independent biological replicates (n = 5), with shaded regions indicating ±1 standard deviation. The middle horizontal line indicates the mean, and the upper and lower horizontal lines represent ±1 standard deviation.

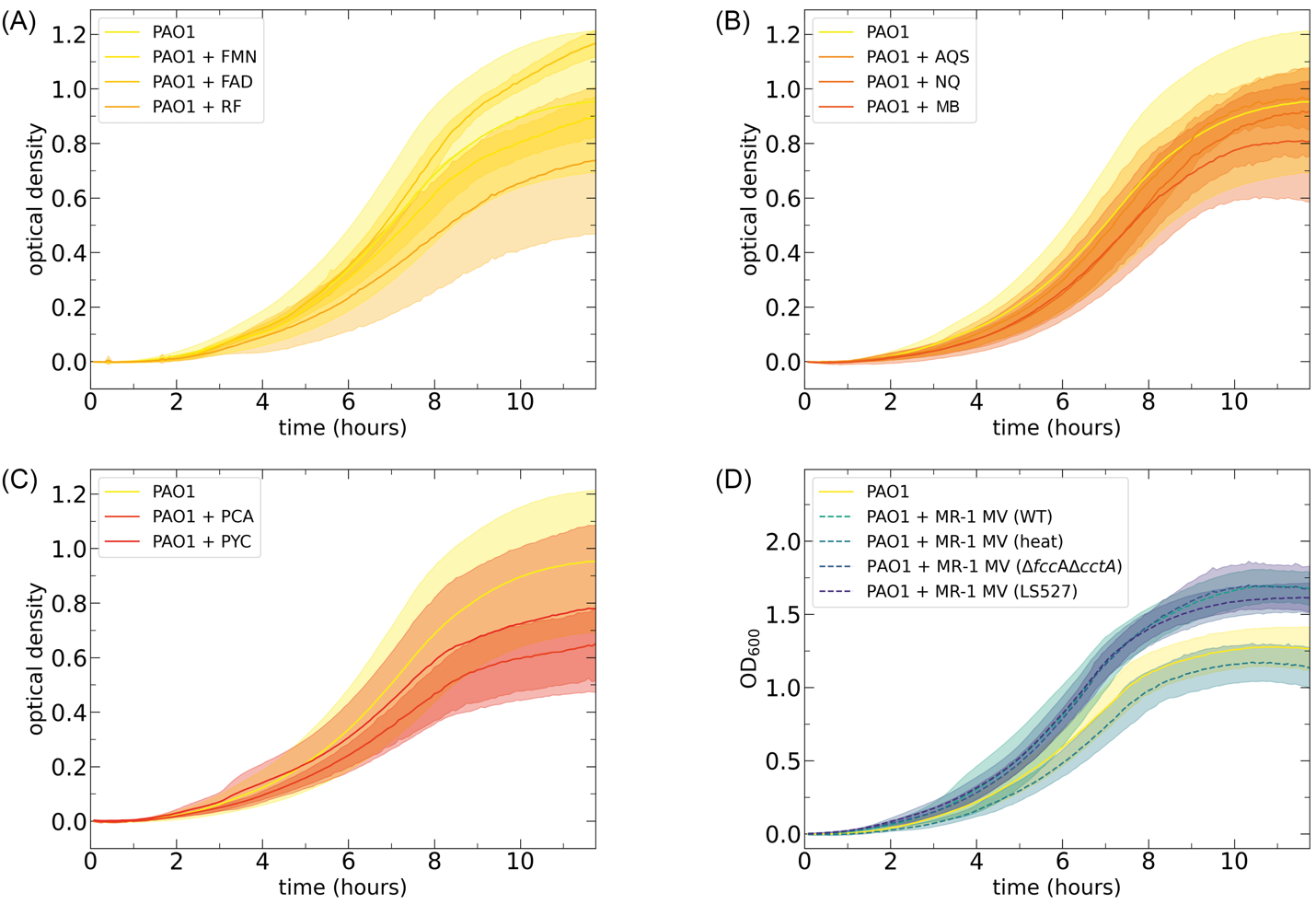

#### Supplementary Figure S8*. Pseudomonas aeruginosa* growth with electron mediators and MV from *Shewanella oneidensis* mutants

Growth curves of *Pseudomonas aeruginosa* in the presence of electron mediators and MV from *S. oneidensis* mutants. We cultured *P. aeruginosa* cells with 100 µM electron mediators or *Shewanella oneidensis* mutants MV (x4 MV concentrations) in LB medium with 100 mM nitrate at 37 ºC in anoxic conditions. The panels show the data (A) with secreted electron mediators from *S. oneidensis*, (B) with artificial electron mediators, (C) with secreted electron mediators from *P. aeruginosa*, and (D) with *S. oneidensis* MV from mutants. FMN: flavin adenine dinucleotide; FAD: flavin mononucleotide; RF: riboflavin; AQS: anthraquinone-2-sulfonate; NQ: 2-hydroxy-1,4-naphthoquinone; MB: methylene blue; PCA: phenazine-1-carboxylic acid; PYC: pyocyanin. Data points represent the mean of independent biological replicates (n ≥ 5), with shaded regions indicating ±1 standard deviation.
